# Epigenetic dysregulation from chromosomal transit in micronuclei

**DOI:** 10.1101/2022.01.12.475944

**Authors:** Albert Agustinus, Ramya Raviram, Bhargavi Dameracharla, Jens Luebeck, Stephanie Stransky, Lorenzo Scipioni, Robert M. Myers, Melody Di Bona, Mercedes Duran, Britta Weigelt, Shira Yomtoubian, Eléonore Toufektchan, Paul S. Mischel, Vivek Mittal, Sohrab Shah, John Maciejowski, Enrico Gratton, Peter Ly, Mathieu F. Bakhoum, Dan Landau, Vineet Bafna, Simone Sidoli, Yael David, Samuel F. Bakhoum

## Abstract

Chromosomal instability (CIN) and epigenetic alterations are characteristics of advanced and metastatic cancers [1-4], yet whether they are mechanistically linked is unknown. Here we show that missegregation of mitotic chromosomes, their sequestration in micronuclei [5, 6], and subsequent micronuclear envelope rupture [7] profoundly disrupt normal histone post-translational modifications (PTMs), a phenomenon conserved across humans and mice as well as cancer and non-transformed cells. Some of the changes to histone PTMs occur due to micronuclear envelope rupture whereas others are inherited from mitotic abnormalities prior to micronucleus formation. Using orthogonal techniques, we show that micronuclei exhibit extensive differences in chromatin accessibility with a strong positional bias between promoters and distal or intergenic regions. Finally, we show that inducing CIN engenders widespread epigenetic dysregulation and that chromosomes which transit in micronuclei experience durable abnormalities in their accessibility long after they have been reincorporated into the primary nucleus. Thus, in addition to genomic copy number alterations, CIN can serve as a vehicle for epigenetic reprogramming and heterogeneity in cancer.

## INTRODUCTION

Chromosomal instability (CIN) drives tumor progression, in part, through the generation of genomic copy number heterogeneity that serves as a substrate for natural selection [2, 4, 8-10]. CIN is associated with metastasis [11], therapeutic resistance [12], and immune evasion [13, 14], and it results from ongoing chromosome missegregation during mitosis [15]. A hallmark of cancer cells with CIN is the presence of lagging chromosomes in anaphase. Missegregating chromosomes frequently end up in micronuclei, whose envelopes are rupture prone, exposing their genomic content to the cytosol [6, 11, 16, 17]. Widespread DNA damage from micronuclear envelope rupture can catalyze genomic abnormalities, including complex chromosomal rearrangements known as chromothripsis [18, 19]. Chromosomes that are encapsulated in micronuclei are often wholly or partially re-integrated into the primary nucleus after the successive mitosis and as such can propagate genetic abnormalities – acquired while in the micronucleus – to daughter cells [18, 19]. While the genomic ramifications of chromosome missegregation has been widely studied, little is known about the impact of chromosomal transit in micronuclei on epigenetic integrity.

## RESULTS

### Abnormal histone posttranslational modifications in micronuclei

To determine the epigenetic consequences of chromosome sequestration in micronuclei, we used high-resolution immunofluorescence microscopy and assessed the status of various canonical histone post-translational modifications (PTMs) in human non-transformed mammary epithelial cells (MCF10A), telomerase-immortalized retinal pigment epithelial cells (RPE1), high-grade serous ovarian cancer (HGSOC) cells (OVCAR-3), as well as human and mouse triple-negative breast cancer (TNBC) cells (MDA-MB-231 and 4T1, respectively). In all five cell lines, we observed substantial differences in the landscape of histone PTMs when comparing primary nuclei and micronuclei. Micronuclei displayed significant reduction in the active transcriptional activating mark, lysine acetylation, at multiple residues along the histone H3 tail (H3K9Ac, H3K14Ac, and H3K27Ac). In addition, the two canonical ubiquitination events on histones, the repressive mono-ubiquitination of histone H2A (H2AK119Ub) and the gene bodies-specific mono-ubiquitination of histone H2B (H2BK120Ub), exhibited substantial reductions, with the latter at a near total loss (**Fig. 1A-B and Supplementary Fig. 1A-B**). This discrepancy in histone PTM patterns between primary nuclei and micronuclei was remarkably conserved among nontransformed and cancer-derived cells as well as across species (**Fig. 1B and Supplementary Fig. 1B**). Interestingly, while certain key methyl-lysine residues on histone H3 were preserved in micronuclei, some, such as H3K9me3, H3K27me3, and H3K36me3, were further enriched, as evidenced by increased fluorescence signal intensity relative to the primary nucleus (**Supplementary Fig. 1C**). Treatment with the pan-histone deacetylase (HDAC) inhibitor, vorinostat, led to near-complete restoration of H3K9Ac, H3K14Ac, and H3K27Ac signals in micronuclei, whereas treatment with the EZH2 inhibitor, GSK126, led to significant reduction in H3K27me3 staining intensity, supporting the specificity of the detected immunofluorescence signal (**Fig. 1C** and **Supplementary Fig. 2A-E**).

**Figure 1:**
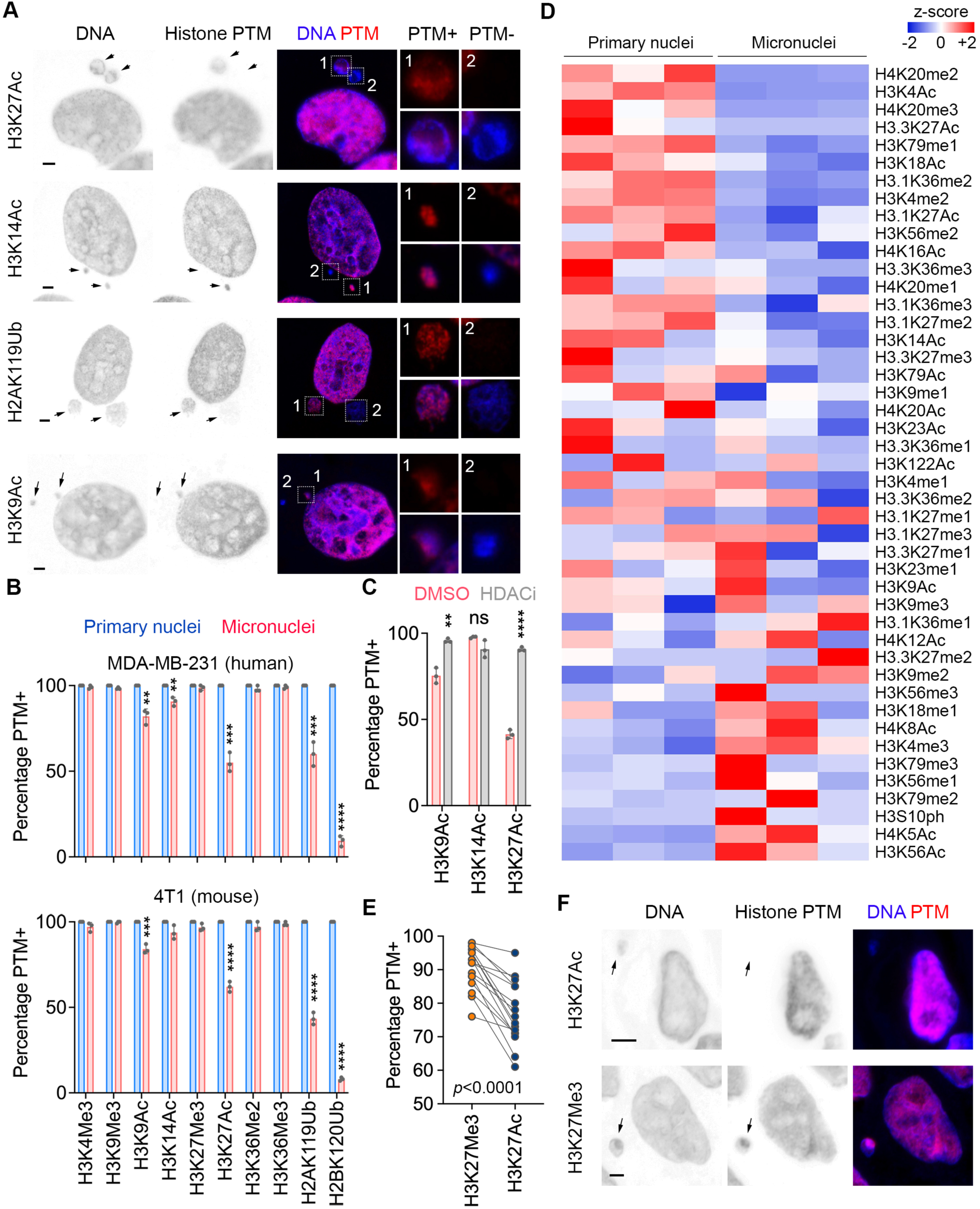
Distinct histone posttranslational modifications in micronuclei. **A**, Representative immunofluorescence images of micronucleated MDA-MB-231 cells stained for DNA (blue) and histone posttranslational modifications (PTMs) (red), arrows point to micronuclei, scale bars 2 μm. **B**, Percentage of primary nuclei and micronuclei with given histone PTMs in MDA-MB-231 and 4T1 cells, ** *p* < 0.01, *** *p* < 0.001, **** *p* < 0.0001, two-sided t-test, n = 3 biological replicates, bars represent mean ± SD. **C**, Percentage of micronuclei with given histone PTMs in MDA-MB-231 cells treated with DMSO or vorinostat (HDAC inhibitor); ** *p* < 0.01, *** *p* < 0.001, **** *p* < 0.0001, two-sided t-test, n = 3 biological replicates, bars represent mean ± SD. **D**, Heat map of z-score of relative abundance of histone PTMs in purified micronuclei and primary nuclei of 4T1 cells, n = 3 biological replicates. **E**, Percentage of micronuclei with H3K27me3 and H3K27Ac in human high grade serous ovarian cancer samples (n = 16), statistical significance tested using two-sided paired *t* test. **F**, Representative immunofluorescence images from HGSOC samples stained for DNA (blue) and either H3K27Me3 or H3K27Ac (red), arrows point to micronuclei, scale bars 2 μm.

To orthogonally validate our microscopy-based findings, we purified and separated primary nuclei and micronuclei from MDA-MB-231 and 4T1 cells by means of selective plasma membrane permeabilization, sucrose gradient ultracentrifugation, followed by fluorescence-activated sorting, as previously described [20] (**Supplementary Fig. 3A**). Immunoblot analysis confirmed loss of both monoubiquitination marks (H2AK119Ub and H2BK120Ub) in micronuclei (**Supplementary Fig. 2F**). Furthermore, unbiased quantitative mass spectrometry survey of canonical histone PTMs confirmed significant reductions in acetylation on multiple lysine residues of histones H3 and H4, compared to primary nuclei (**Fig. 1D and Supplementary Fig. 3B**). Overall, the histone marks between primary nuclei and micronuclei were markedly distinct, following a nearly mutually exclusive pattern (**Fig. 1D**). These changes could not be attributed to differences in the total levels of histone H3 between primary nuclei and micronuclei (LC-MS/MS AUC ± s.e.m. = 3.05×10^10^ ± 8.85×10^9^ and 3.39×10^10^ ± 1.36×10^10^ for primary nuclei and micronuclei, respectively, p = 0.85, two-sided t-test) or specific degradation of the histone tail as the explanation for the differential enrichment of the histone marks, some of which were differentially enriched on the same lysine residue (e.g., loss of acetylation and gain of trimethylation on H3K27).

To examine whether the same trends exist in human tumors, we stained 16 HGSOC samples for two key histone marks (H3K27me3 and H3K27Ac). In line with our findings in cell lines, we observed relative preservation of the H3K27me3 mark in micronuclei compared with a more widespread loss of H3K27Ac in each sample (**Fig. 1E-F**). Together, our data indicate that micronuclei exhibit conserved changes in histone PTMs which include preservation of stable heterochromatin-associated histone marks (e.g., H3K27me3) and loss of the more dynamic PTMs that denote transcriptionally active chromatin, such as H3K27Ac and H2BK120Ub.

### Mitotic errors and micronuclear envelope rupture alter histone PTMs

We reasoned that the observed abnormalities in histone PTMs might result from micronuclear envelope rupture [6, 7], which provides aberrant cytosolic access to the chromatin [7]. To test this, we co-stained histone PTMs with cGAS, a cytosolic double-stranded DNA sensor that binds genomic DNA in ruptured micronuclei [11, 16, 17, 21]. Our results indicate that micronuclei lacking histone H3 acetylation (K9Ac, K14Ac, and K27Ac) as well H2AK119Ub were much more likely to display cGAS colocalization, suggesting that these marks are lost upon micronuclear envelope rupture (**Fig. 2A-B**). This observation was consistent across human HGSOC samples, where a significant loss of H3K27Ac was selectively seen in ruptured compared with intact micronuclei (**Fig. 2C-D**). On the sample-level, the fraction of micronuclei with cGAS staining was strongly correlated with the percentage of micronuclei lacking H3K27Ac (**Supplementary Fig. 4A**). Importantly, treatment of MDA-MB-231 cells with the HDAC inhibitor, vorinostat, restored histone H3 lysine acetylation in ruptured micronuclei, and to a much lesser extent in the intact ones (**Fig. 2E**).

**Figure 2:**
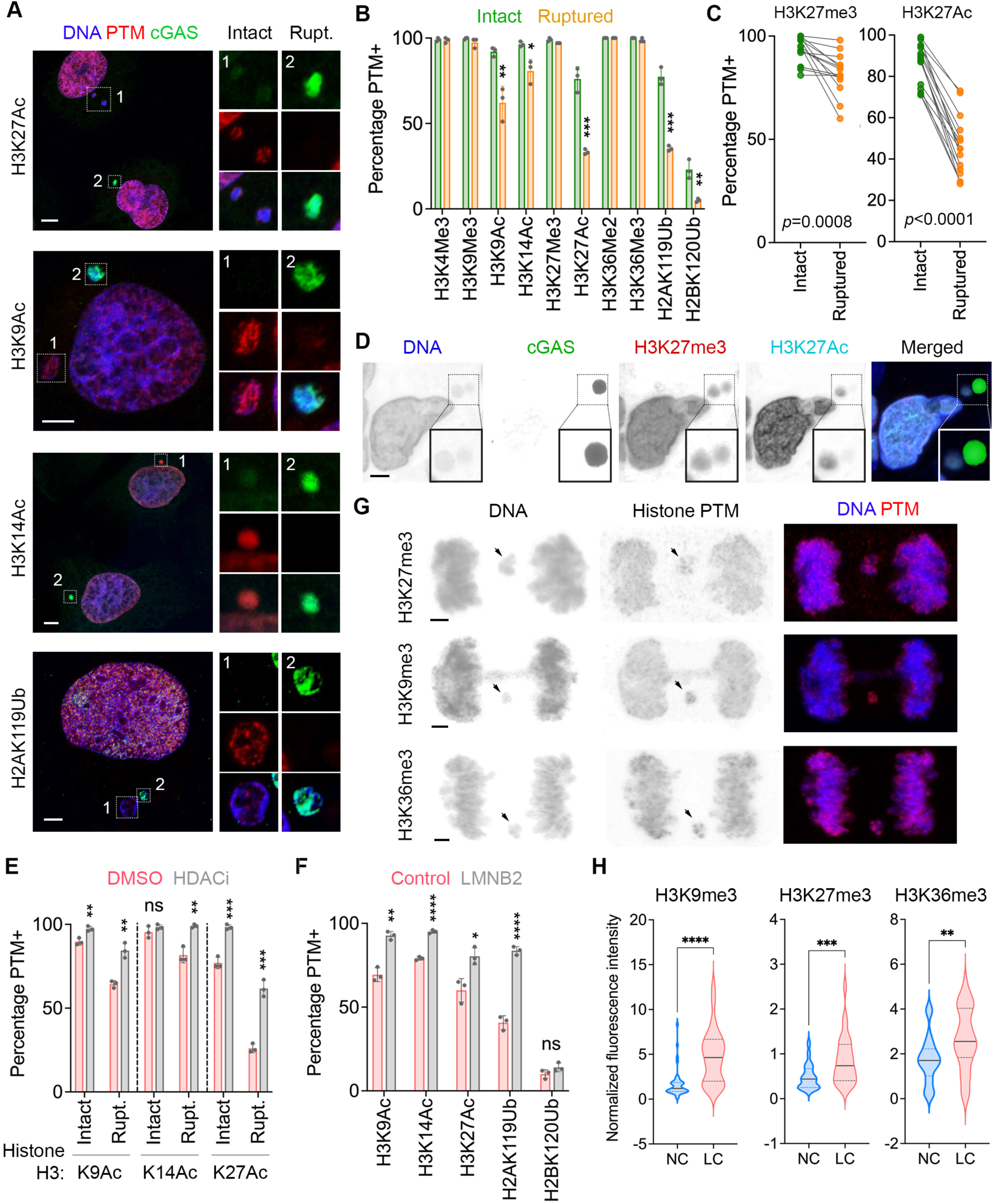
Micronuclear rupture and chromosome missegregation alter histone PTMs. **A**, Representative immunofluorescence images of MDA-MB-231 cells with micronuclei stained for DNA (blue), histone PTMs (red), and cGAS (green), scale bars 5 μm. **B**, Percentage of intact and ruptured micronuclei with given histone PTMs from MDA-MB-231 cells; * *p* < 0.05, ** *p* < 0.01, *** *p* < 0.001, two-sided t-test, n = 3 biological replicates, bars represent mean ± SD. **C**, Percentage of intact and ruptured micronuclei with H3K27me3 and H3K27Ac in human HGSOC (n = 16), statistical significance tested using two-sided paired *t* test. **D**, Representative immunofluorescence images from HGSOC samples stained for DNA (blue), cGAS (green in), H3K27Me3 (red), or H3K27Ac (cyan), scale bar 2 μm. **E**, Percentage of intact and ruptured micronuclei with given histone PTMs in MDA-MB-231 cells treated with DMSO or vorinostat (HDAC inhibitor); ns, not significant, ** *p* < 0.01, *** *p* < 0.001, two-sided t-test, n = 3 biological replicates, bars represent mean ± SD. **F**, Percentage of micronuclei with given histone PTMs in control and lamin B2-overexpressing (LMNB2) MDA-MB-231 cells, * *p* < 0.05, ** *p* < 0.01, **** *p* < 0.0001, two-sided t-test, n = 3 biological replicates, bars represent mean ± SD. **G**, Representative immunofluorescence images of MDA-MB-231 cells undergoing anaphase with lagging chromosomes (denoted by arrows). Cells were stained for DNA (blue) and histone PTMs (red), scale bars 2 μm. **H**, Violin plots showing the normalized fluorescence intensity distribution of histone PTMs on normal chromosomes (NC) and lagging chromosomes (LC) during mitosis, n > 28 cells; ** *p* < 0.01, *** *p* < 0.001, **** *p* < 0.0001, two-sided Mann-Whitney test, solid and dashed bars in the plot represent the median and quartiles, respectively.

In contrast, the H2BK120Ub mark was lost in nearly all micronuclei, whereas many stable heterochromatin-associated marks (H3K9me3, H3K27me3, and H3K36me3) were enriched, irrespective of micronuclear rupture status, suggesting rupture-independent changes (**Fig. 2B** and **Supplementary Fig. 4B-C**).

To directly test the contribution of micronuclear rupture to changes in histone PTMs, we overexpressed Lamin B2, which suppresses micronuclear envelope collapse [7]. As expected, Lamin B2 overexpression in MDA-MB-231 cells significantly reduced the fraction of micronuclei with cGAS staining (**Supplementary Fig. 4D-F**). It also selectively rescued abnormalities in rupture-associated histone PTMs (H3K9Ac, H3K14Ac, H3K27Ac, and H2AK119Ub), with no impact on rupture-unassociated changes such as loss of H2BK120Ub or enrichment of lysine trimethyl marks on Histone H3 (**Fig. 2F and Supplementary Fig. 4G**).

We then asked whether alterations in histone PTMs that were not associated with micronuclear rupture might arise during the prior mitosis and are subsequently inherited with the lagging chromosome, thus predating the formation of the micronucleus. We examined the fluorescence intensity of 10 canonical histone PTMs during anaphase, comparing lagging chromosomes with the remaining normally segregating chromosomes. All surveyed histone PTMs were present on lagging chromosomes except for H2BK120Ub, which was completely absent in anaphase (**Fig. 2G and Supplementary Fig. 5A-B**). Interestingly, the relative fluorescence intensities of H3K9me3, H3k27me3, and H3K36me3 – all of which were enriched in micronuclei – were also higher on lagging chromosomes compared to their normally segregating counterparts (**Fig. 2G-H**).

Unlike all the other examined canonical histone marks, H2BK120Ub was conspicuously absent from mitotic cells, as previously reported [22], from prometaphase until early telophase (**Supplementary Fig. 5B**). While chromosomes in the primary nucleus quickly regained this mark upon late telophase, micronuclei did not (**Supplementary Fig. 5B**). We reasoned that the unequal redeposition of this mark might be due to the biased subcellular localization of the known H2BK120 ubiquitin ligases (RNF20/40) near the spindle poles [23]. This would favor direct encapsulation of RNF20/40 into the primary nucleus during telophase, as opposed to the micronucleus that arises mostly from chromosomes that lag near the spindle midzone where RNF20/40 are also depleted [6, 23]. To test this, we sought to promote the formation of micronuclei arising from chromosomes that missegregate all the while remaining near the spindle poles without transiting through the midzone by inhibiting CENP-E. CENP-E is a kinesin-like motor protein that promotes chromosome congression to the metaphase plate [24]. Treatment of MDA-MB-231 cells with the CENP-E-specific inhibitor, GSK923295 [25], led to a significant increase in micronuclei containing H2BK120Ub (**Supplementary Fig. 5C**). Collectively, these data suggest that some histone marks are disrupted as a direct result of micronuclear envelope rupture whereas others arise from the malpositioning of lagging chromosomes during anaphase, predating the formation of the micronucleus and persisting with the chromosome into the subsequent cell cycle.

### Altered chromatin accessibility in micronuclei

To test how abnormalities in the histone PTM landscape might alter chromatin structure in micronuclei, we used three orthogonal approaches starting with fluorescence lifetime imaging microscopy (FLIM), which provides information regarding chromatin states at nanoscale spatial resolutions [26]. Using FLIM, we reliably identified euchromatic and heterochromatic regions in primary nuclei (**Fig. 3A-B**). Expectedly, chromatin compaction in intact micronuclei resembled that of heterochromatic regions (**Fig. 3A-B**). On the other hand, ruptured micronuclei displayed a wide distribution of chromatin compaction states, ranging from extremely compact to open chromatin (**Fig. 3A-B**). Interestingly, even within individual ruptured micronuclei there was marked spatial heterogeneity as indicated by diverse regions of chromatin accessibility (**Fig. 3A**, *right*), which we reasoned might be due to partially digested DNA. Using a second approach, we treated MDA-MB-231 cells with a transposase loaded with fluorophore-tagged adaptors to image chromatin accessibility at subcellular resolutions, a technique known as ATAC-see [27]. Fluorescence signal, which arises from primers installed by the transposase, was readily apparent in primary nuclei and markedly diminished in micronuclei (**Fig. 3C-D** and **Supplementary Fig. 6A**). Treatment with either EZH2 or HDAC inhibitors led to a significant increase in the ATAC-see signal in micronuclei to levels equivalent to primary nuclei (**Fig. 3D**), suggesting that enrichment for H3K27me3 along with the loss of H3K27Ac among other marks leads to an overall reduction in chromatin accessibility in micronuclei.

**Figure 3:**
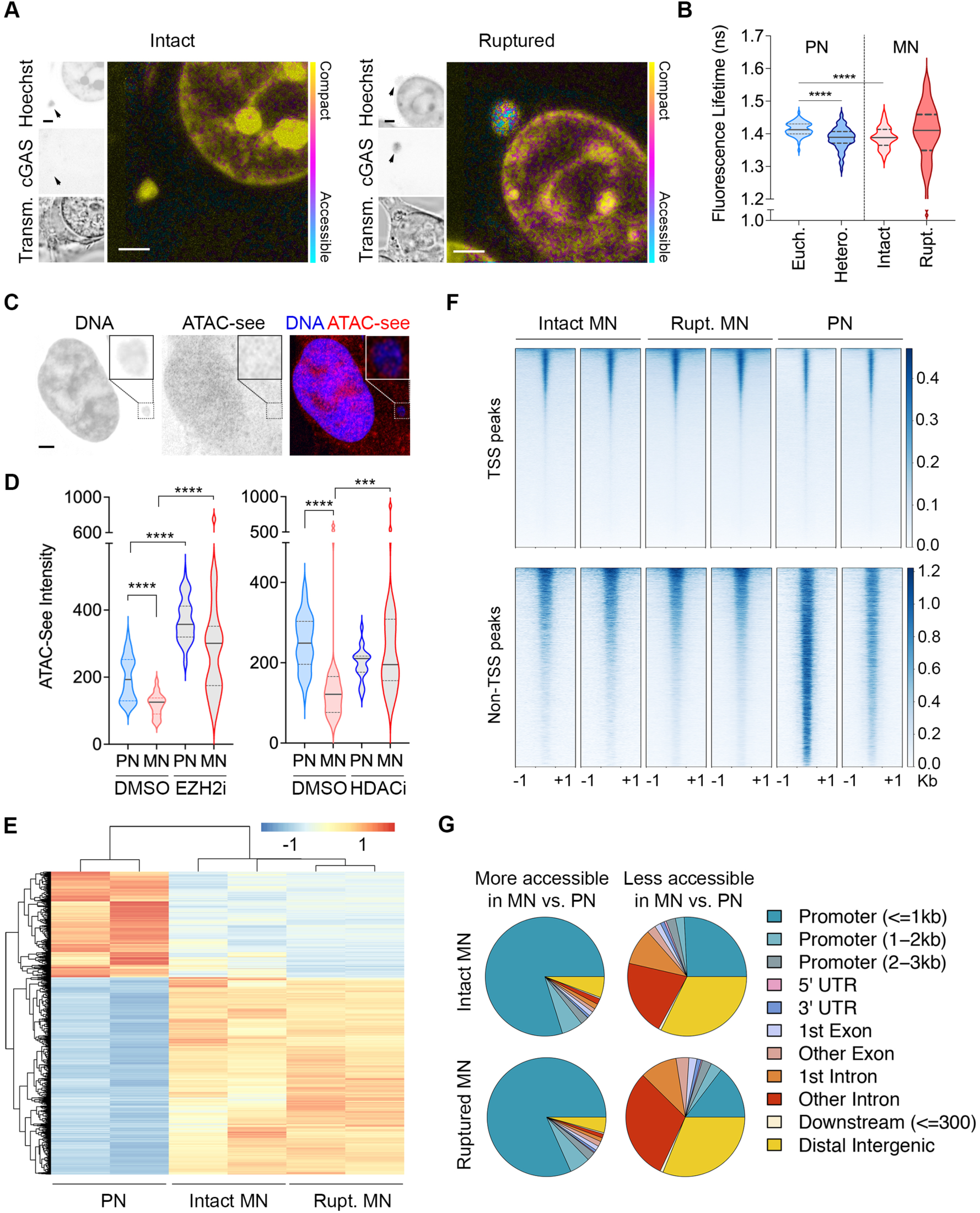
Altered chromatin accessibility in micronuclei. **A**, Representative fluorescence-lifetime imaging microscopy images of cGAS-GFP expressing 4T1 cells with micronuclei stained with Hoechst. Pseudo-color panels represent Hoechst intensity lifetime, as a readout for chromatin compactness. **B**, Violin plots representing the distribution of fluorescence lifetime in euchromatic and heterochromatic regions of primary nuclei, as well as ruptured and intact micronuclei, **** *p* < 0.0001, n = 73 cells (31 intact and 42 ruptured micronuclei) from five biological replicates, two-sided Mann-Whitney test, solid and dashed bars in the plot represent the median and quartiles, respectively. **C**, Representative ATAC-see fluorescence images of MDA-MB-231 cells with micronuclei stained for DNA (blue), scale bar 2 μm. **D**, Violin plots representing the distribution of ATAC-see signal intensity quantification of primary nuclei (PN) and micronuclei (MN) of MDA-MB-231 cells treated with either vehicle control (DMSO), GSK126, an EZH2 inhibitor (EZH2i), or Vorinostat, an HDAC inhibitor (HDACi), *** *p* < 0.001, **** *p* < 0.0001, two-sided Mann-Whitney test, n = 5 biological replicates, solid and dashed bars in the plot represent the median and quartiles, respectively. Statistical significance tested using. **E**, Heat map showing differentially accessible genomic peaks from isolated primary nuclei (PN), intact (Intact MN) and ruptured (Rupt. MN) micronuclei from 4T1 cells, n = 2 biological replicates. **F**, Heat map representing chromatin accessibility at transcription start sites (TSS peaks) and all other sites (non-TSS peaks) in primary nuclei (PN), intact (Intact MN) and ruptured (Rupt. MN) micronuclei from 4T1 cells, n = 2 biological replicates. **G**, Pie charts representing the regions that are either more (left charts) or less (right charts) accessible in either intact (top) or ruptured (bottom) micronuclei compared to primary nuclei in 4T1 cells, n = 2 biological replicates.

Using a third and orthogonal approach, we performed a classical ATAC-seq analysis on primary nuclei, intact micronuclei, and ruptured micronuclei, purified from H2B-mCherry and cGAS-GFP expressing 4T1 cells by sorting micronuclei based on their GFP signal after sucrose gradient ultracentrifugation [20] (**Supplementary Fig. 6B-D**). Principal component analysis (PCA) based on merged peaks across the samples (n = 14,676) indicated differences between intact, ruptured, and primary nuclei (**Supplementary Fig. 7A**). Micronuclei and primary nuclei exhibited a significant number of differentially accessible peaks (**Fig. 3E**). To our surprise, this differential accessibility involved a profound positional bias; while many transcriptional start sites (TSS) were more accessible in micronuclei, non-TSS peaks were significantly less accessible (**Fig. 3F**). Indeed, a more in-depth positional analysis revealed an over-whelming positional bias in micronuclei towards more accessible promoters compared with reduced accessibility within intronic and distal intergenic regions (**Fig. 3G**).

Furthermore, the accessibility of the gene bodies mirrored the differential accessibility of their associated promoters (**Supplementary Fig. 7B-C**). Finally, there was moderate concordance in promoter accessibility between intact and ruptured micronuclei yet, with a subset that was selectively associated with the rupture status (**Supplementary Fig. 7D**). Furthermore, the accessibility of the gene bodies mirrored the differential accessibility of their associated promoters (**Supplementary Fig. 7B-C**). Finally, there was moderate concordance in promoter accessibility between intact and ruptured micronuclei yet, with a subset that was selectively associated with the rupture status (**Supplementary Fig. 7D**).

We next asked if genes that were more accessible in micronuclei were transcriptionally enriched in human cancers with CIN. Gene-set enrichment analysis (GSEA) comparing human breast tumors belonging to the top and bottom tertiles of the fraction of genome altered (FGA) according to The Cancer Genome Atlas (TCGA) revealed significant positive transcriptional enrichment of micronuclei-accessible genes in FGA^high^ tumors (**Supplementary Fig. 7E**). Conversely, transcripts of genes that were less accessible in micronuclei were correspondingly depleted in FGA^high^ relative to FGA^low^ tumors (**Supplementary Fig. 7E**). Collectively, our data demonstrate that micronuclei have an overall more compact chromatin structure with a marked positional bias that favors accessible chromatin near some promoter regions and heterochromatin at intronic and distal intergenic regions.

### CIN as a catalyst for lasting epigenetic dysregulation

To test whether CIN and micronuclear envelope rupture can lead to global epigenetic and transcriptional dysregulation, we subjected control or Lamin B2 overexpressing chromosomally stable TP53-knocout RPE-1 cells to long-term treatment with reversine, an MPS1 inhibitor known to induce chromosome segregation defects [28], at doses well below its IC50 concentrations (**Supplementary Fig. 8A-B**). Reversine-treated cells exhibited an increase in anaphase chromosome missegregation, micronuclei formation, and pervasive numerical and structural chromosomal aberrations indicative of CIN (**Supplementary Fig. 8C-E**). Long-term induction of CIN with reversine led to significant changes in both chromatin accessibility as well as the overall transcription (**Fig. 4A-B**). While Lamin B2 on its own did not significantly impact either, it partially diminished some of the epigenetic and transcriptional changes induced by reversine (**Fig. 4A-C**). PCA based on all differentially accessible genomic peaks, RNA expression of all genes, or the expression of the subset of genes that were differentially accessible in micronuclei, indicated differences between control and Lamin B2 overexpressing cells only when treated with reversine but not with DMSO (**Fig. 4A-C and Supplementary Fig. 8F**). In line with our observations in micronuclei, there was increased accessibility at promoter sites in reversine-treated cells, which was significantly reduced upon lamin B2 overexpression (**Supplementary Fig. 8G**). Importantly, genes that were overall more accessible in micronuclei were also transcriptionally enriched in reversine-treated cells, whereas those which were exclusively accessible in ruptured micronuclei were correspondingly downregulated upon Lamin B2 overexpression, in line with reduced micronuclear rupture (**Fig. 4D**). Thus, inducing CIN promotes global changes in chromatin accessibility and transcription that mirrors patterns of chromatin alterations seen in micronuclei.

**Figure 4:**
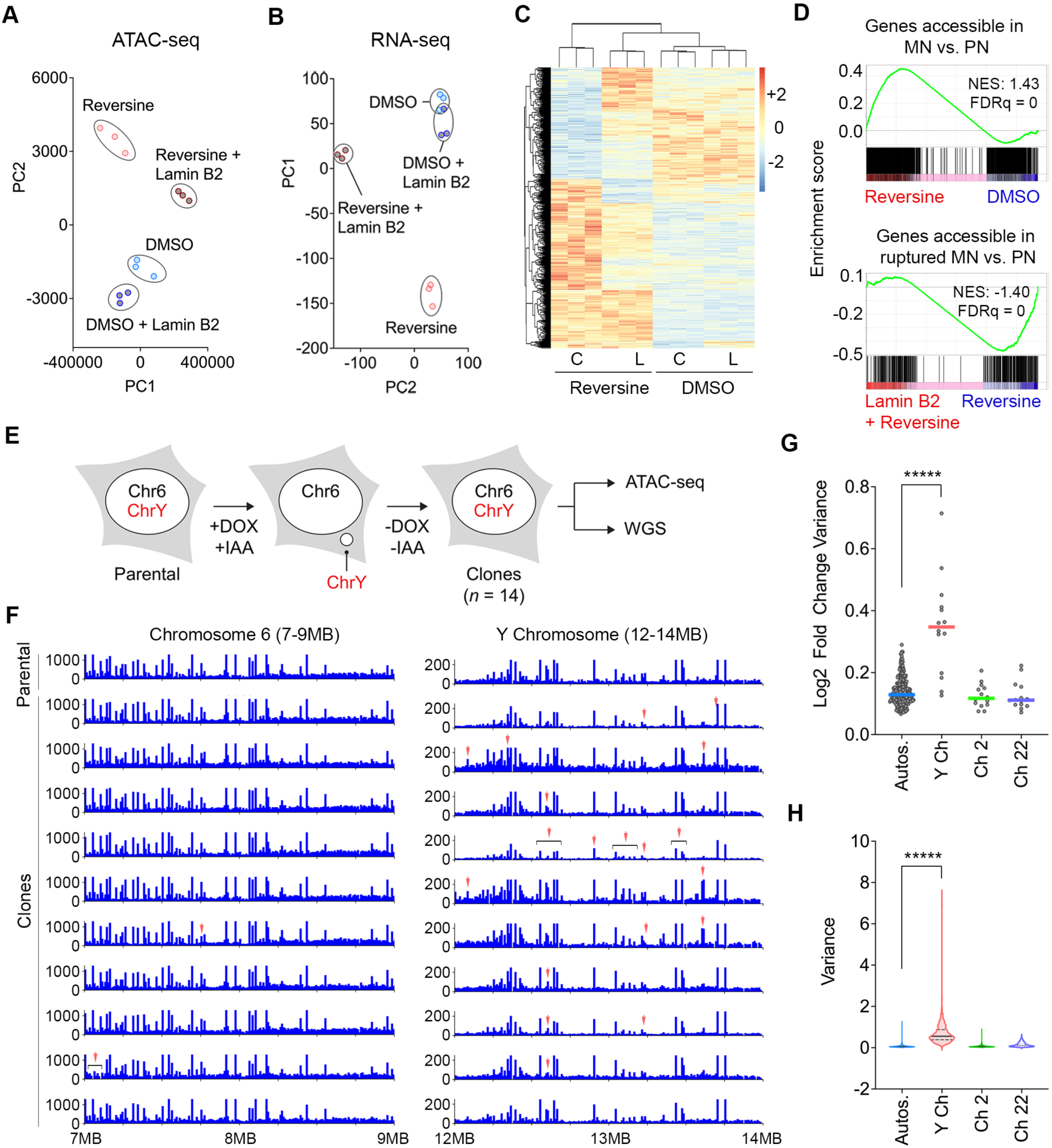
Chromosomal instability drives long-term changes in chromatin accessibility. **A-B**, Principal component analysis (PCA) plot of TP53 KO hTERT-RPE1 cells based on accessibility from ATAC-seq (a) or RNA expression obtained from RNA-seq (b), n = 3 biological triplicates. **C**, Heat map representing chromatin accessibility in control (C) or Lamin B2 (L) overexpressing TP53 KO hTERT RPE-1 cells treated with reversine or vehicle control (DMSO), n = 3 biological replicates. **D**, Enrichment plots comparing the expression of genes whose promoters are accessible in all micronuclei (top) comparing DMSO or reversine treated RPE-1 cells (top), enrichment plot comparing the expression of genes whose promoters are accessible exclusively in ruptured micronuclei comparing control or Lamin B2 overexpressing RPE-1 cells both treated with reversin (bottom). **E**, Experimental schema depicting CEN-SELECT system used in DLD-1 cells to select clones that have exclusively mis-segregate and micronucleate the Y chromosome and subsequently reincorporated it to the nucleus in the next cell cycle. Y Ch = Y chromosome, Ch7 = chromosome 7, an example of any autosomes. **F**, Genome viewer plot showing copy number-normalized signals in a 2KB region of chromosome 6 (as a control auto-some) and the Y chromosome. Top row represents parental cells which did not undergo missegregation, while the rest represent single cell clones that underwent missegregation and transient micronucleation of the Y chromosome. Red arrow denotes differentially accessible peaks compared to parental cells. **G**, Interclonal comparison of fold change variance of ATAC-seq peaks in 14 DLD-1 single clones showing the Y chromosome and chromosomes 2 and 22 as randomly selected autosomes. Autos = autosomes, Y ch = Y chromosome, Ch 2 = chromosome 22, Ch 22 = chromosome 22. Bar represents median. ***** *p* < 0.00001, two-sided Mann-Whitney test. **H**, Violin plots representing intraclonal variance across 10kb segments in each of 14 DLD-1 clones. Autos = autosomes, Y ch = Y chromosome, Ch 2 = chromosome 22, Ch 22 = chromosome 22. Distribution represents 86,666 calculated values for autosomes, 730 calculated values for Y chromosome. ***** *p* < 0.00001, Mann-Whitney test.

To directly ask whether CIN-induced epigenetic abnormalities are selective to chromosomes that transit in micronuclei, we used an inducible Y chromosome-specific missegregation system established in otherwise chromosomally stable DLD-1 colorectal cancer cells [18, 29]. This approach allows efficient doxycycline- and auxin-inducible micronucleation of the Y chromosome caused by centromere inactivation without affecting the autosomes or X chromosome [29]. The addition of a Y-encoded selection marker enabled the derivation of single cell-derived clones in which the Y chromosome has been re-incorporated into the primary nucleus after transient micronucleation [18], allowing us to assess the long-term epigenetic consequences that are specific to the Y chromosome in each clone as compared to the remaining autosomes (**Fig. 4E**). To determine the long-term impact of transient micronucleation events on chromatin accessibility, we performed ATAC-seq on the parental cells with an intact Y chromosome and 14 clones harboring Y chromosomes that were transiently induced into micronuclei and subsequently re-incorporated into the primary nucleus [18]. ATAC-seq reads were then normalized to DNA copy number obtained from whole-genome sequencing of each clone [18] (**Supplementary Fig. 9** and **Methods**). Relative to parental cells, we observed striking interclonal variability in the copy number-corrected accessibility profile on the Y chromosome but not the autosomes or X chromosome (**Fig. 4F-G and Supplementary Fig. 10A**). Irrespective of chromosome size or any underlying genomic clonal copy number alterations, the accessibility patterns of the autosomes and X chromosome were largely maintained. In contrast, the Y chromosome showed extreme fold-change of accessibility that was weakly correlated with the extent of its genomic copy number alteration (**Supplementary Fig. 10B**). In line with our findings from comparing the different clones, the variance in accessibility along 10kb genomic segments within individual clones on the Y chromosome also far exceeded that of the autosomes and X chromosome (**Fig. 4H, Extended Data Figs. 9 and 10C**). Thus, chromosomes that transit in micronuclei are subject to long-lasting epigenetic dysregulation even after reincorporation into the primary nucleus.

## DISCUSSION

Prior research has mostly focused on the genomic impact of chromosomal transit in micronuclei [6, 18, 19, 30] and little is known about the epigenetic integrity of unstable chromosomes. Our work demonstrates that the transient sequestration of chromosomes in micronuclei has profound consequences on chromatin organization and can disrupt epigenetic states long after the initial micronucleation event when the chromosome is later re-integrated into the primary nucleus (**Fig. 4E-H**). The overall compact chromatin organization in micronuclei is likely driven by the relative enrichment of stable heterochromatin-associated histone marks and the loss of the more dynamic histone PTMs such as H3K27Ac (**Fig. 1**). Nonetheless, the surprising positional accessibility bias seen in micronuclei near promoter regions suggests that micronuclei might catalyze chromatin reorganization and the aberrant transcription of genes that are otherwise silenced, in line with our observations in experimental systems and human breast tumors (**Fig. 4D and Supplementary Fig. 7E**).

Epigenetic reprogramming is a key driver of tumorigenesis and recent studies have identified numerous alterations in epigenetic modifying enzymes as a source for abnormalities seen in cancer [3, 31-36]. Given the stochastic nature of chromosome encapsulation in micronuclei, we propose that CIN can also drive epigenetic reprogramming by altering the chromatin organization of entire chromosomes in the span of a single cell cycle. This has the potential to promote cell-to-cell variability in genomic accessibility patterns as well as gene expression, serving as a substrate for natural selection. CIN-induced epigenetic reprogramming might also act as means for dosage compensation [37] to buffer potentially deleterious chromosome copy number alterations.

Large-scale changes among histone PTMs in cytosol-accessible chromatin may also act as a biochemical signal impacting downstream processes such as senescence, the sensing of cytosolic double-stranded DNA by the cGAS-STING pathway, as well as DNA repair in micronuclei. For instance, H3K36me3 has been proposed to determine the choice of DNA repair pathways [38, 39], and its enrichment in micronuclei would likely favor the patching of pulverized chromosomes [5, 6] via non-homologous end joining, leading to patterns observed during complex rearrangements and chromothripsis [18, 19].

Finally, the remarkable conservation of histone PTMs in micronuclei across species and in non-transformed cells, along with the positional bias in micronuclear chromatin reorganization, has potentially important implications to processes involved in cancer as well as normal physiology. For instance, this work raises the tantalizing possibility that micronuclei might serve as catalysts for oncogene amplification during tumorigenesis. The increased accessibility near promoter regions in the setting of rampant chromosomal breaks in micronuclei [6] form the substrates needed for the formation of circular extrachromosomal DNA, which has an overall accessible chromatin state and a high density of oncogenes [30, 40, 41]. In non-transformed settings, there is also evidence that changes in histone PTMs in micronuclei might play a physiological role.

Loss of histone H3 acetylation in the germline micronucleus of the unicellular eukaryote, Tetrahymena, has been proposed as a mechanism for selective transcriptional silencing in the setting of increased replication [42], which is required for sexual reproduction. Thus, epigenetic alterations in micronuclei likely represent an evolutionary conserved phenomenon that is aberrantly co-opted by cancer cells.

## Acknowledgements

We would like to thank members of the Bakhoum and David Laboratories (MSKCC), Daniel Bronder (MSKCC), Ashley Laughney (Weill Cornell Medicine), Neil Vasan and Benjamin Izar (Columbia University Medical Center) for constructive feedback. We would also like to thank the MSKCC genomics, molecular cytology, flow cytometry, and molecular cytogenetic cores. Work in the Bakhoum laboratory is supported by the High-Risk High-Reward Program from the Office Of The Director, National Institutes Of Health (DP5OD026395), the National Cancer Institute (Breast Cancer SPORE P50CA247749 and R01CA256188-01), the Department of Defense Congressionally Directed Medical Research Program, the Burroughs Wellcome Fund Career Award for Medical Scientists, the Parker Institute for Immunotherapy at MSKCC, the STARR Cancer Consortium, the Josie Robertson Foundation, and the MSKCC core grant (P30-CA008748). Work in the David laboratory is supported by the Josie Robertson Foundation, the Pershing Square Sohn Cancer Research Alliance, the NIH (CCSG core grant P30 CA008748, MSK SPORE P50 CA192937 and R35 GM138386), the Parker Institute for Cancer Immunotherapy (PICI), and the Anna Fuller Trust. In addition, the David lab is supported by Mr. William H. Goodwin and Mrs. Alice Goodwin and the Commonwealth Foundation for Cancer Research and the Center for Experimental Therapeutics at MSKCC. The Sidoli lab gratefully acknowledges the Leukemia Research Foundation (Hollis Brownstein New Investigator Research Grant), AFAR (Sagol Network GerOmics award), Deerfield (Xseed award), Relay Therapeutics, Merck and the NIH Office of the Director (1S10OD030286-01). AA is supported by the PhRMA Foundation Predoctoral Fellowship. VB is supported in part by grants U24CA264379, R01GM114362 from the NIH. RMM is supported by the Medical Scientist Training Program grant from the NIGMS of the NIH under award number T32GM007739 to the Weill Cornell/Rockefeller/Sloan Kettering Tri-institutional MD-PhD program.

## METHODS

### Cell culture

Cell lines (MDA-MB-231, 4T1, RPE-1) were purchased from American Type Culture Collection (ATCC). MCF10A p53 KO and RPE-1 p53KO were kind gifts from the Maciejowski lab at MSKCC. MDA-MB-231 and RPE-1 were cultured in DMEM (Corning) while 4T1 was cultured in RPMI (Corning), both media were supplemented with 10% FBS and 50 U/mL penicillin/streptomycin. MCF10A were cultured in DMEM/F-12 (Corning) supplemented with 5% horse serum, 100 ng/mL cholera toxin, 10 μg/mL insulin, 20 ng/mL human epidermal growth factor, and 0.5 μg/mL hydrocortisone. All cells were periodically tested for mycoplasma.

### Immunofluorescence microscopy and histone PTM quantification

Cells were grown on coverslip until 80-90% confluent. Cell fixation was performed using ice-cold (−20°C) methanol for 15 minutes in the freezer. Cells were then permeabilized with 0.5% triton in TBS-BSA 1% solution for 5 minutes at room temperature. Subsequently, cells were washed with TBS-BSA 1% and incubated for 5 minutes at room temperature. Antibodies (see **Supplementary Table 1** for complete list and dilutions) were diluted with 1%BSA-TBS solution and incubated in 4°C overnight. Then, DAPI (0.5 μg/mL) was added together with secondary antibodies diluted in TBS-BSA 1%. Cells were mounted with Fluoro-Gel with tris buffer (Electron Microscopy Sciences). Images were collected as described later. Histone PTM positive/negative quantification was done in triplicates for each histone PTMs, with 100 micronuclei counted per replicate. Histone PTM intensity quantification was done using ZEN 2 (blue edition) software, where the fluorescence intensity of the corresponding PTM is normalized to its DAPI channel fluorescence intensity.

### Immunoblots

*Western Blots*: Cells were lysed with 1x RIPA Buffer (EMD Millipore), kept on ice and vortexed 3 times in 10 minutes interval. The lysates were then centrifuged at 10,000 rpm for 10 minutes and debris were discarded. Protein was quantified using DC protein assay (Bio-Rad) and an equal amount of protein are loaded in each SDS-PAGE gel well and ran at 120 V for 1 hour. Proteins were then transferred to a nitrocellulose membrane using standard protocol in Bio-Rad Trans-Blot Turbo transfer system. Membrane was blocked using TBS blocking buffer (Li-Cor) incubated with primary antibody listed in Supplementary Table 2 overnight at 4 C. Secondary antibody incubation was done for 1 hour at room temperature. Scanning was done using the Odyssey CLx system (Li-Cor). *Dot Blots*: Isolated nuclei were lysed with 1x RIPA Buffer (EMD Millipore), kept on ice and vortexed 3 times in 10 minutes interval. The lysates were then centrifuged at 10,000 rpm for 10 minutes and debris were discarded. No quantification was done for dot blot, an approximation of 1:20 ratio between micronuclei and primary nuclei was used instead. A 2 μL of the RIPA extract of either micronuclei or primary nuclei fraction are blotted on top of nitrocellulose membrane and air dried for at least 1 hour. Membrane was then blocked using TBS blocking buffer (Li-Cor) incubated with primary antibody listed in Supplementary Table 2 overnight at 4 C. Secondary antibody incubation was done for 1 hour at room temperature. Scanning was done using the Odyssey CLx system (Li-Cor) and quantification was done using ImageStudio software (Li-Cor). Antibodies used for immunoblots are listed in **Supplementary Table 2**.

### Drug treatments

Reversine (Cayman Chemical Company) was given to cells at the concentration of 0.5 μM for either 24 hours (for MDA-MB-231) or 48 hours (4T1). SAHA (Sigma-Aldrich), HDAC inhibitor was used at the concentration of 0.5 μM for 24 hours, and GSK-126 (Xcess-Bio), EZH2 methyltransferase inhibitor was given to cells at the concentration of 5 μM for 5 days. Long term reversine treatment was done 24 hours after cells were passaged, and media was replaced with normal cell media 48 hours after. GSK923295 (Selleck Chemicals) was used at the concentration of 50 nM for 24 hours. DMSO was used as vehicle control.

### Plasmids and molecular biology

H2B-mCherry and GFP-cGAS expressions were achieved using plasmids obtained from the Maciejowski Lab. Puromycin at the concentration of 2 μg/mL was used to select for cells expressing both proteins, and subsequently, fluorescence-activated cell sorting was used to enrich for cells expressing the respective proteins. LaminB2-mCherry overexpression was achieved using plasmid obtained from the Hetzer laboratory. Blasticidin was used at the concentration of 10 μg/mL to select for cells overexpressing LaminB2.

### Fluorescence lifetime imaging microscopy (FLIM)

*Microscopy*: Fluorescence lifetime images were collected with a Zeiss LSM 880 (Zeiss, Thornwood, NY) microscope equipped with a Spectra-Physics MaiTai HP laser (Spectra-Physics, Santa Clara, CA) for 2-photon excitation, tuned at 800 nm, and collected using a Zeiss 63x/1.41 NA Oil objective. The fluorescence signal was collected by a photomultiplier tube (H7422P-40, Hamamatsu, Bridgewater, NJ), and recorded using a FLIMbox (Model A320 ISS, Champaign, IL) to obtain the lifetime information. The pixel dwell time for the acquisitions was 16 μs and the images were taken with sizes of 256×256 pixels accumulating 15 frames. The data from each pixel were recorded and analyzed using the SimFCS software (Laboratory for Fluorescence Dynamics, University of California, Irvine, CA) to perform the phasor transformation and with custom Matlab code to quantify the lifetime. The phasor position was corrected for the instrument response function (IRF) by acquiring the fluorescence of a solution with known lifetime (0.4 mM Coumarin 6 dissolved in Ethanol, τ = 2.5 ns). The lifetime was determined as τ=s/ωg, where τ is the fluorescence lifetime, (g,s) are the phasor coordinated and ω is the laser frequency (80 MHz). *Cell labeling and image processing*: the samples were labelled with one drop per well of Hoechst 33342 (NucBlue, Thermofisher Scientific) and imaged after 20 minutes from incubation. For every field of view for FLIM, we also recorded a 2-channel fluorescence intensity image corresponding to the Hoechst 33342 and EGFP emissions to identify the ruptured micronuclei. The four subregions considered (heterochromatin, euchromatin, intact and ruptured micronuclei) were manually determined according to the following criteria: 1) Heterochromatin: On the Hoechst 33342 channel, bright spots in the nucleus correspond to heterochromatin enriched regions in NIH-3T3 cells. 2) Euchromatin: On the Hoechst 33342 channel, regions in the nucleus with low Hoechst 33342 signal were classified as Euchromatin-enriched regions. 3) Micronuclei: On the Hoechst 33342 channel, micronuclei were identified as nucleic acids positive compartments with smaller size compared to the main nucleus. They were further classified as: intact, where no signal in the EGFP channel or ruptured, where positive signal in the EGFP channel. The subregions selected above were used to segment the lifetime image and a single value of lifetime was obtained as the median value of the entire subregion. For every cell, one value of lifetime for heterochromatin, euchromatin and micronuclei (either ruptured or intact) was obtained, for a total of 73 cells (31 intact, 42 ruptured micronuclei) over five biological replicas. The values were cleared from outliers by the MATLAB function “isoutlier”.

### ATAC-see with immunofluorescence

Tn5 production, transposome assembly, and ATAC-See were performed as described by Chen, et al [27] with some adjustments for subsequent immunofluorescence. Briefly, cells were grown on coverslip until 80-90% confluent, and were fixed using 1% PFA in PBS at room temperature for 10 minutes. Permeabilization was done in lysis buffer (10 mM Tris–Cl, pH 7.4, 10 mM NaCl, 3 mM MgCl2, 0.01% Igepal CA-630) for 10 minutes at room temperature. Washing was done using PBS, twice for 5 minutes each at room temperature. The transposase mixture solution (25 μL 2× TD buffer, final concentration of 100 nM Tn5-ATTO-59ON, adding dH2O up to 50 μL) was added on the coverslip and incubated for 30 min at 37 °C in a humid chamber box. Coverslips were then washed three times with PBS containing 0.01% SDS and 50 mM EDTA for 15 minutes at 55 °C each time. After washing, coverslips were subjected to the immunofluorescence procedure described before, starting from washing with TBS-BSA 1%.

### Super-resolution immunofluorescence microscopy on cultured cells

Images were acquired with an inverted Zeiss LSM880 (Carl Zeiss Microscopy GmbH, Germany) equipped with an Airy Scan detector (gain 850, digital gain 1) in super resolution mode. The 848 × 848 pixels images were acquired with a Plan-Apochromat 63x/1.4 Oil objective, using a pixel size of 40 nm and a dwell time of 4.96 ms. The optical configuration used split the laser lines (405nm at 1% power, 488nm at 3% and 561nm at 7%) with a 488/561/633 MBS and an invisible light beam splitter at 405. In detection we applied a band pass filter 495-550 nm and a long pass 570 nm. Each image is acquired as a Z-stack of 5 stacks, with an interval of 500 nm; a maximum intensity projection and an Airyscan processing with default parameters were performed for all the images before quantification using the Zeiss ZEN black software. For the ATAC-see images, only two laser lines were used, 405 nm and 594 nm, at 1% and 30% power respectively, using the same optical configuration described above.

### Micronuclei purification

The micronuclei purification was performed as described previously [43], with the exception that primary nuclei fraction is also collected. Briefly, cells were expanded in 245 × 245 × 25 mm dishes and treated with reversine as specified previously. Cells were then harvested and washed DMEM. Washed cells were resuspended in pre-warmed (37°C) DMEM supplemented with 10 μg/ml cytochalasin B (Sigma-Aldrich) at a concentration of around 5×10^6^ cells/mL DMEM and incubated at 37°C for at least 30 minutes. Subsequently, cells were centrifuged at 300g for 5 minutes and resuspended in cold lysis buffer (10 mM Tris-HCl, 2 mM Mg-acetate, 3 mM CaCl2, 0.32 M sucrose, 0.1 mM EDTA, 0.1% (v/v) NP-40, pH 8.5) freshly complemented (with 1 mM dithiothreitol, 0.15 mM spermine, 0.75 mM spermidine, 10 μg/ml cytochalasin B and protease inhibitors) at a concentration of 10^7^ cells/ml lysis buffer. Resuspended cells were then dounce-homogenized 10 times using a loose-fitting pestle. Then, lysates were mixed with an equal volume of cold 1.8 M sucrose buffer (10 mM Tris-HCl, 1.8 M sucrose, 5 mM Mg-acetate, 0.1 mM EDTA, pH 8.0) freshly complemented (with 1 mM dithiothreitol, 0.3% BSA, 0.15 mM spermine, 0.75 mM spermidine) before use. In a 50 mL conical tube, 10 mL of the mixture was then layered on top of a two-layer sucrose gradient (20 mL of 1.8 M sucrose buffer on top of 15 mL 1.6 M sucrose buffer). This mixture was then centrifuged in a swing bucket centrifuge at 950g for 20 min at 4°C. The first resulting 2 mL top fraction is discarded; next 5–6 mL mostly contain micronuclei is collected, the next 3 mL mostly containing primary nuclei is also collected in a separate container. Collected fractions were diluted 1:5 with FACS buffer (ice cold PBS freshly supplemented with 0.3% BSA, 0.1% NP-40 and protease inhibitors). Diluted MN were then centrifuged at 950g in JS-5.2 swing bucket centrifuge for 20 min at 4°C. The resulting supernatant was removed by aspiration and either micronuclei or primary nuclei was resuspended in 2–4 mL of FACS buffer supplemented with 2 μg/ml DAPI (however, no DAPI was used for micronuclei purification for ATAC-seq experiments). Resuspended samples were filtered through a 40 μm ministrainer (PluriSelect) into FACS tubes. MN were then sorted by FACSAria (BD Biosciences) into FACS buffer at the MSKCC Flow Cytometry Core Facility. Default FSC and DAPI thresholds were lowered and a log scale was used to visualize MN, for ATAC-seq micronuclei purification, FSC and mCherry were used instead of FSC and DAPI. Sorted MN were centrifuged at 4000g in a swing bucket rotor for 20 min at 4°C and the pellets were stored at −80°C for later use, or processed immediately if they were to be used for ATAC-seq.

### RNA-Seq

RNA was extracted using the protocols in the Qiagen RNeasy mini plus kit (Qiagen). Library preparations are TruSeq stranded, and sequenced using PolyA RNA-sequencing workflow, with read depth of 80-100 million reads, 100 bp paired end reads. Library preparation and sequencing were performed in Memorial Sloan Kettering Cancer Center Integrated Genomics Core (IGO). RNAseq data were analyzed using Basepair toolkit (https://www.basepairtech.com/) using the expression count (STAR) workflow to obtain FPKM values. Principle Component Analysis (PCA) was computed using randomized principal component analysis applied to the normalized count matrix of all genes (Fig. 4B) or genes that were differentially accessible between micronuclei and primary nuclei (Supplementary Fig. 8F) using Partek Flow software, version 10.0 (Partek Inc). Gene set enrichment analysis (GSEA) was performed on normalized gene counts from control and Lamin B2 overexpressing TP53 KO hTERT-RPE1 cells treated with reversine or DMSO (Fig. 4D) using the GSEA software (The Broad Institute), with the following parameters: metric for ranking genes, Signal2Noise; weighted enrichment statistics; permutation type, gene_set. The gene sets used in GSEA were derived from ATAC-seq analysis of chromatin accessibility in intact or ruptured micronuclei in 4T1 (Supplementary Fig. 7E). Mouse gene symbols were converted to human gene symbols using the Ensembl biomaRt package (version 2.48.3). Expression datasets from the TCGA breast cancer cohort were accessed through cbioportal[44] (https://www.cbi-oportal.org/study/summary?id=brca_tcga_pan_can_at-las_2018). GSEA comparing tumors belonging to the top or bottom tertile of the fraction of genome altered, as reported by TCGA, was performed as described above, and using the same parameters as above (Supplementary Fig. 7F-G).

### ATAC-Seq

The ATAC-seq procedure on cells were done in accordance to Buenrostro et al [45]. The ATAC-seq procedure on nuclei samples was done immediately after isolation using the same protocol without the lysis step. The ratio of primary nuclei to micronuclei was 1:20. Libraries were sequenced with 50 bp paired end reads to approximately 50 million reads per sample for cells, whereas for nuclei samples, sequencing was done with 50 bp paired end reads to 350 million total reads for a triplicate of each sample at the Memorial Sloan Kettering Cancer Center Integrated Genomics Core (IGO). For DLD-1 ATAC-Seq experiments, the raw fast data were analyzed using Basepair toolkit (https://www.basepairtech.com/). The raw reads were trimmed using fastp to remove low-quality bases from reads (quality < 20) and adapter sequences. The trimmed reads were aligned using Bowtie2 [46] to UCSC genome assembly (hg38). (RAMYA for MN ATAC-Seq). Reads were mapped to the hg38 (RPE-1) or mm10 (MCF10A) genome assembly using bowtie2 using the follow parameters: -X2000 --no-mixed --no-discordant and reads were filtered using samtools for the 1804 FLAG and mapq score of 30. PCR duplicates were identified using the MarkDuplicates function from Picard tools. Peaks were called per sample using macs2 with the following parameters: callpeak -f BAM -g hs --nomodel --shift 37 --extsize 73 --keep-dup all -B --SPMR --call-summits -q 1e-2. Final peaks across each dataset was determined based on 250bp extension from the summit of each peak and merged across all samples using bedtools. Normalized bigwig files were generated using the bamCoverage function from deeptools using RPKM normalization. Heatmaps for genomic regions were generated using the deeptools computeMatrix scale-region and plotHeatmap functions. Differential peak calling was performed using DESeq2 on raw counts for the merged peaks (an adjusted p-value cut-off of 0.01 and log2FC of 1.5 was applied to the mouse MCF10A cell lines and an adjusted p-value of 0.05 and log2FC of 0.75 was applied to the RPE-1 cell line). The normalized counts for peaks were used to plot the heatmaps of differential peaks. Pie charts were generated using the plotAnnoPie function in the ChIPseeker R package.

### ATAC-Seq normalization for DLD-1 cells

#### CNV calling from WGS data

WGS CRAM files were obtained from EGA (dataset ID EGAD00001004163), and unmapped from GRCh37 using samtools (version 1.3.1), and then aligned to GRCh38 using BWA-MEM (version 0.7.17-r1188). Copy number profiles were called using CNVKit (version 0.9.7). Integer copy numbers were obtained using the CNVKit “call” command, and non-integer copy numbers were obtained from the log2 CN ratios reported in the CNVKit .cns files using PrepareAA’s convert_cns_to_bed.py script (https://github.com/jluebeck/PrepareAA). *ATAC-seq Data Filtering*: ATAC-seq alignments were filtered from the BAM files to remove PCR duplicates, reads aligning with mitochondrial DNA, and alignments with low mapping quality (MAPQ score < 15). *Signal Calculation*: We used the ATAC-seq signal not to identify peaks corresponding to active transcriptional sites but to measure the total accessibility of DNA within a region. We generated ATAC-seq read counts for 10000 bp genomic windows using samtools (version 1.13). We then calculated the signal values for each window by normalizing the read counts by copy number, chromosome length, and total atac reads across all chromosomes. Windows overlapping with blacklisted regions, or with a copy number close to 0 (< 0.3) were excluded from further analysis.

The normalized signal of a clone c in a given window i with total atac reads T[i] and copy number C[i] was calculated as

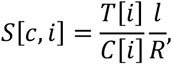

where R is the total number of atac reads mapped across all chromosomes in clone c; l denotes the ‘effective length’ of the chromosome as follows. Let P denote the expected ploidy (P=1 for chr Y; P=2 otherwise) of the chromosome, and l[i],C[i] denote the length (in bp) and average copy number, respectively, of window i in the chromosome. Then,

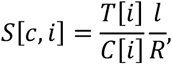

In the case of the Y chromosome, we only used the region up to 26.68 mbp for the above calculations because the Y chromosome did not have any confidently aligned ATAC reads beyond 26.68 Mbp until the telomeric region in any of the clones (including the parental clone).

### Differential Analysis of Y chromosome accessibility

The normalized ATAC signals of a clone can be compared to the parental signals by taking the log signal fold-changes with respect to the parental signal in the same window. The signal fold-change of a clone c in a given window i is given by:

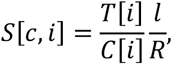

where p stands for the parental clone. *Intra-clone variability and change in accessibility*: we computed the mean and variance of log signal fold changes for each chromosome in a single clone to understand the global changes in accessibility. To control for high fold-changes caused by error in low read count windows, windows where neither the clone nor the parent has greater than 200 reads were excluded from this calculation. *Inter-clone variability*: to understand whether the Y chromosome accessibility changes were similar for different clones, we calculated the variance of log signal fold changes of different clones in the same genomic window. Windows where no clone or parent has greater than 200 reads were excluded. In addition, only clones with a copy number > 0.3 in the window were included for the variance calculation. *Calculating Extreme fold-changes in chrY*: we checked for the presence of extreme signal fold changes in Y chromosome in comparison to autosomes by computing the mean and standard deviation of all autosomal fold changes and computing the percentage of chromosome Y windows in each clone that have a fold change that is more than 5 standard deviations away from the mean. To see the correlation between shattering of Y chromosome and extreme fold changes, we calculated the Pearson rank correlation p-value for the percent of windows with extreme fold changes and the fraction of chromosome with a copy number equal to the expected copy number. In addition to the genome-based normalization method described above, we have repeated the entire analysis of DLD-1 cells using a chromosome-based normalization method and obtained similar results (see **Supplementary Methods**). The motivation for this alternative normalization is based on the irregular distribution of total ATAC-seq reads per chromosome among the clones.

### Immunofluorescence staining of human samples

The Immunofluorescence detections of cGas, H3K27Me3 and H3K27Ac were performed at Molecular Cytology Core Facility of Memorial Sloan Kettering Cancer Center using Discovery Ultra processor (Ventana Medical Systems.Roche-AZ). After 32 min of heat and CC1 (Cell Conditioning 1, Ventana cat#950-500) retrieval, the tissue sections were blocked first for 30 min in Background Blocking reagent (Innovex, catalog#: NB306). A mouse monoclonal cGAS: LSBio (Cat#: LS C757990) was used in 1:200 dilution. The incubation with the primary antibody was done for 5 hours, followed by biotinylated anti-mouse secondary (Vector Labs, MOM Kit BMK-2202) in 5.75ug/mL. Blocker D, Streptavidin-HRP and TSA Alexa488 (Life Tech, cat#B40932) prepared according to the manufacturer’s instruction in 1:150 for 16 min. A mouse monoclonal H3K27Me3 (Active Motif, Cat#: 61017) was used in 1:500 dilution. The incubation with the primary antibody was done for 6 hours, followed by biotinylated anti-mouse secondary (Vector Labs, MOM Kit BMK-2202) in 5.75ug/mL. Blocker D, Streptavidin-HRP and Tyramide-CF594 (Biotium, cat.#92174) were prepared according to manufacturer instruction in 1:2000 for 16 min. A rabbit polyclonal H3K27Ac: Abcam (Cat#: ab4729) was used in 0.05 ug/ml concentration. The incubation with the primary antibody was done for 6 hours, followed by 60 minutes incubation with biotinylated goat anti-rabbit IgG (Vector labs, cat#:PK6101) in 5.75ug/ml. Blocker D, Streptavidin-HRP and TSA Alexa 647 (Life Tech, cat#B40958) is prepared according to the manufacturer’s instruction in 1:150 for 16 min. All slides were counterstained in 5ug/mL DAPI [dihydrochloride(2-(4-Amidinophenyl)-6-indolecarbamidine dihydrochloride], Sigma D9542, for 5 minutes at room temperature, mounted with anti-fade mounting medium Mowiol [Mowiol 4-88 (CALBIOCHEM code: 475904)] and coverslipped. The 956 × 956 pixels Z-stacks images were acquired with an inverted Zeiss LSM880, using the same optical configuration previously described. The acquisition was performed with pixel size and dwell time of 40 nm and 4.39 ms respectively, and using the following lasers and relative powers: 405 nm at 0.4%, 488nm at 3%, 561 nm at 0.5%, and 647nm at 0.5%. The images were processed as previously described, using the Zeiss ZEN black software.

### Histone extraction and digestion for mass spectrometry

Histone proteins were extracted from the cell pellet as described[47] to ensure good-quality identification and quantification of single histone marks. Briefly, histones were acid-extracted with chilled 0.2 M sulfuric acid (5:1, sulfuric acid : pellet) and incubated with constant rotation for 4 h at 4°C, followed by precipitation with 33% trichloroacetic acid (TCA) overnight at 4°C. Then, the supernatant was removed and the tubes were rinsed with ice-cold acetone containing 0.1% HCl, centrifuged and rinsed again using 100% ice-cold acetone. After the final centrifugation, the supernatant was discarded and the pellet was dried using a vacuum centrifuge. The pellet was dissolved in 50 mM ammonium bicarbonate, pH 8.0, and histones were subjected to derivatization using 5 μL of propionic anhydride and 14 μL of ammonium hydroxide (all Sigma Aldrich) to balance the pH at 8.0. The mixture was incubated for 15 min and the procedure was repeated. Histones were then digested with 1 μg of sequencing grade trypsin (Promega) diluted in 50mM ammonium bicarbonate (1:20, enzyme:sample) overnight at room temperature. Derivatization reaction was repeated to derivatize peptide N-termini. The samples were dried in a vacuum centrifuge. Prior to mass spectrometry analysis, samples were desalted using a 96-well plate filter (Orochem) packed with 1 mg of Oasis HLB C-18 resin (Waters). Briefly, the samples were resuspended in 100 μl of 0.1% TFA and loaded onto the HLB resin, which was previously equilibrated using 100 μl of the same buffer. After washing with 100 μl of 0.1% TFA, the samples were eluted with a buffer containing 70 μl of 60% acetonitrile and 0.1% TFA and then dried in a vacuum centrifuge.

### Liquid Chromatography MS/MS Acquisition and data analysis

Samples were resuspended in 10 μl of 0.1% TFA and loaded onto a Dionex RSLC Ultimate 300 (Thermo Scientific), coupled online with an Orbitrap Fusion Lumos (Thermo Scientific). Chromatographic separation was performed with a two-column system, consisting of a C-18 trap cartridge (300 μm ID, 5 mm length) and a picofrit analytical column (75 μm ID, 25 cm length) packed in-house with reversed-phase Repro-Sil Pur C18-AQ 3 μm resin. Histone peptides were separated using a 30 min gradient from 1-30% buffer B (buffer A: 0.1% formic acid, buffer B: 80% acetonitrile + 0.1% formic acid) at a flow rate of 300 nl/min. The mass spectrometer was set to acquire spectra in a data-independent acquisition (DIA) mode. Briefly, the full MS scan was set to 300-1100 m/z in the orbitrap with a resolution of 120,000 (at 200 m/z) and an AGC target of 5×10e5. MS/MS was performed in the orbitrap with sequential isolation windows of 50 m/z with an AGC target of 2×10e5 and an HCD collision energy of 30. Histone peptides raw files were imported into EpiProfile 2.0 software [48]. From the extracted ion chromatogram, the area under the curve was obtained and used to estimate the abundance of each peptide. In order to achieve the relative abundance of post-translational modifications (PTMs), the sum of all different modified forms of a histone peptide was considered as 100% and the area of the particular peptide was divided by the total area for that histone peptide in all of its modified forms. The relative ratio of two isobaric forms was estimated by averaging the ratio for each fragment ion with different mass between the two species. The resulting peptide lists generated by EpiProfile were exported to Microsoft Excel and further processed for a detailed analysis.

## SUPPLEMENTARY FIGURES

**Supplementary Figure 1:**
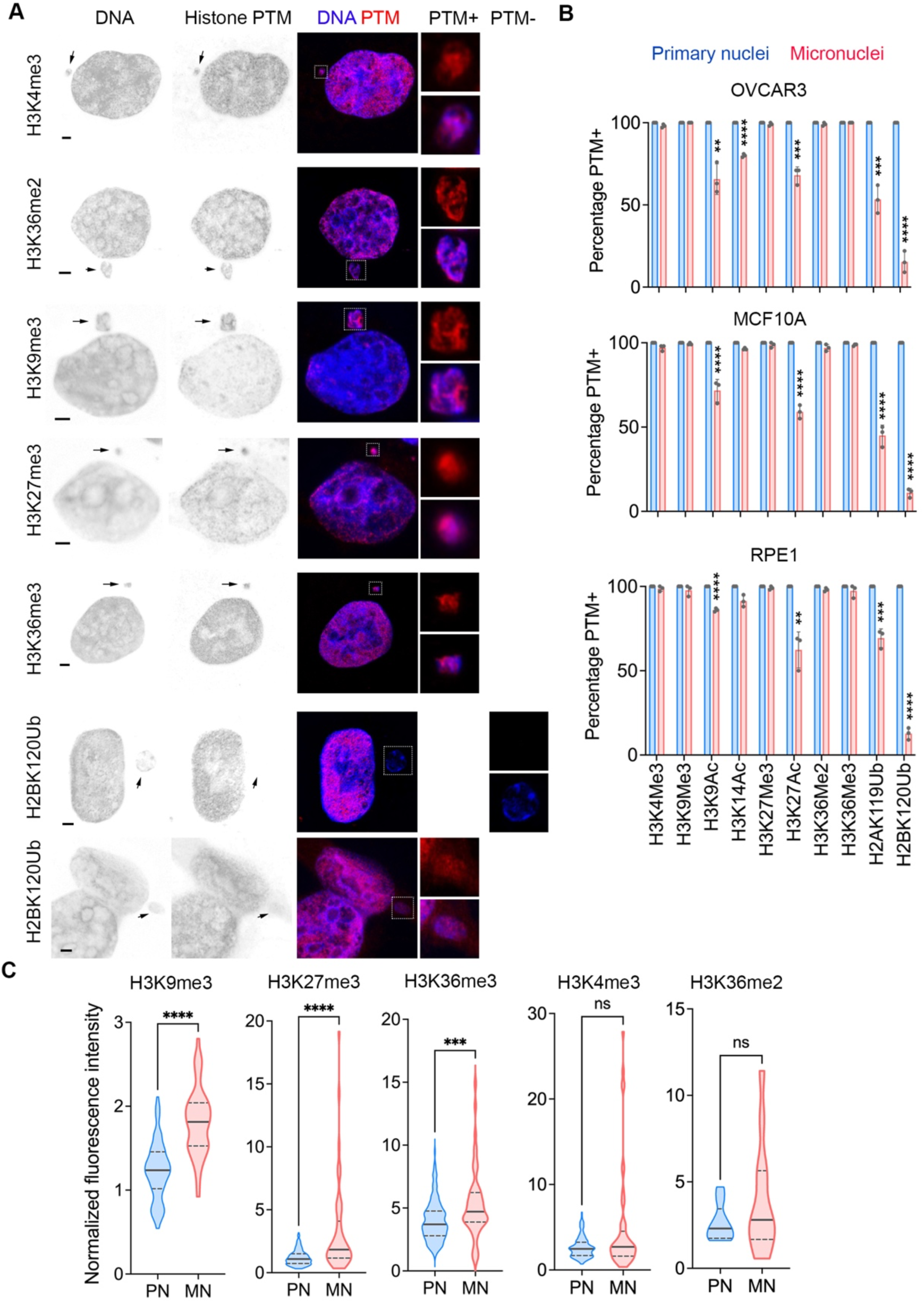
Micronuclei have distinct histone PTM profiles compared with primary nuclei. **A**, Representative immunofluorescence images of micronucleated MDA-MB-231 cells stained for DNA (blue) and histone posttranslational modifications (PTMs) (red), arrows point to micronuclei, scale bars 2 μm. **B**, Percentage of primary nuclei and micronuclei with given histone PTMs in OVCAR3, MCF10A, and RPE-1 cells, ** *p* < 0.01, *** *p* < 0.001, **** *p* < 0.0001, two-sided t-test, n = 3 biological replicates, bars represent mean ± SD. **C**, Violin plot showing the normalized fluorescence intensity distribution of histone PTMs in 13 micronuclei (MN) and accompanying primary nuclei (PN) for H3K36Me2, n > 50 MN and accompanying PN in the same field of view for other PTMs; ns not significant, *** *p* < 0.001, **** *p* < 0.0001, two-sided Mann-Whitney test. Solid and dashed lines in the plot represent the median and quartiles, respectively.

**Supplementary Figure 2:**
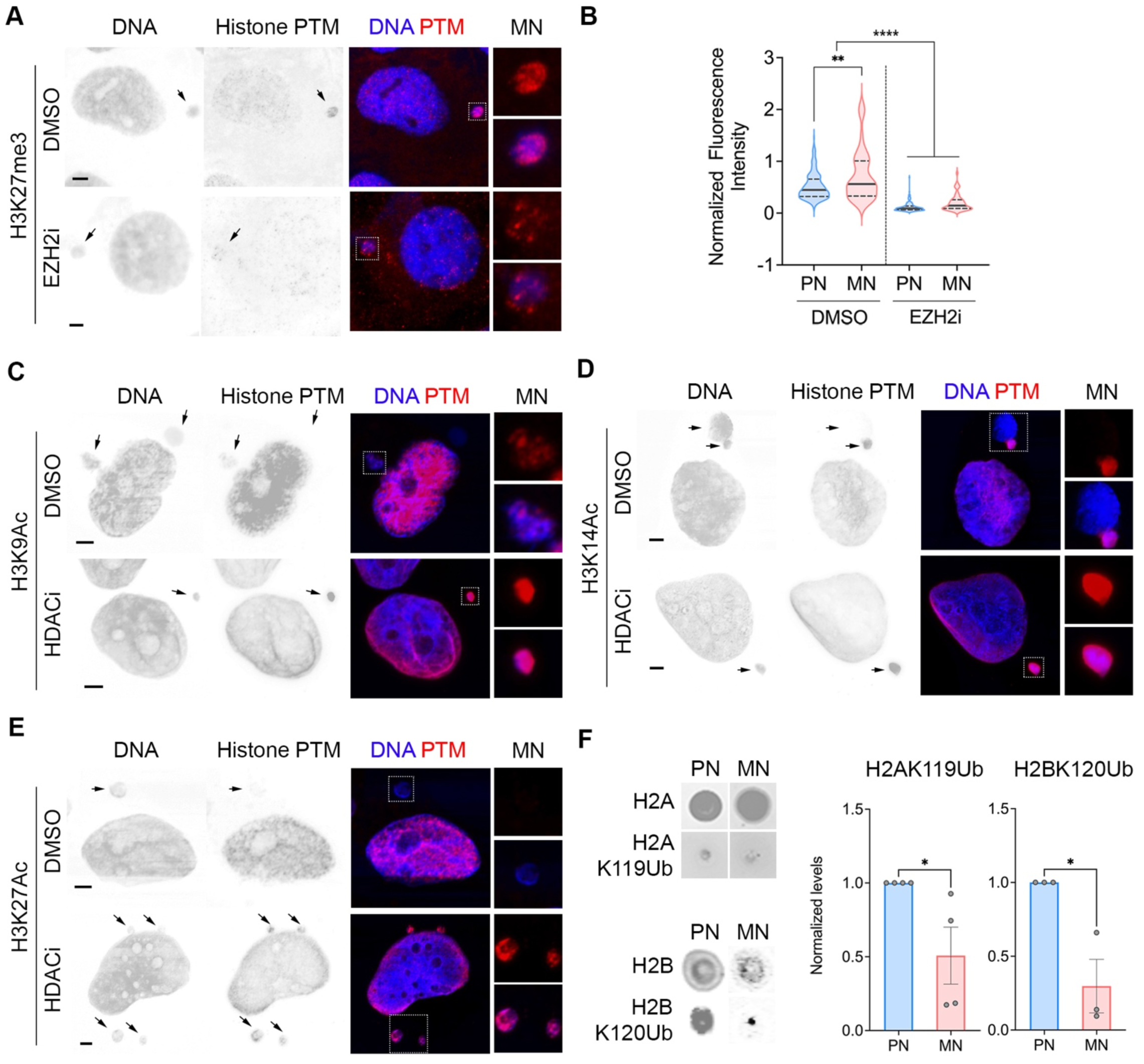
Epigenetic modifying drugs alter histone PTMs in micronuclei. **A**, Representative immunofluorescence images of micronucleated cells stained for DNA (blue) and histone PTMs (red) treated with vehicle control (DMSO) or GSK126 EZH2 inhibitor (EZH2i), arrows point to micronuclei, scale bars 2 μm. **B**, Violin plot showing normalized fluorescence intensity distribution of H3K27Me3 from immunofluorescence experiment in MDA-MB-231, n > 50 micronuclei (MN) and the accompanying primary nuclei (PN). Cells were either treated with vehicle control (DMSO) or GSK126 EZH2 inhibitor (EZH2i); ** *p* < 0.01, **** *p* < 0.0001, two-sided Mann-Whitney test. Solid and dashed lines in the plot represent the median and quartiles, respectively. **C-E**, Representative immunofluorescence images of micronucleated cells stained for DNA (blue) and histone PTMs (red) treated with vehicle control (DMSO) or vorinostat, an HDAC Inhibitor (HDACi). Arrows point to micronuclei, scale bars 2 μm. **F**, Dot blots of H2AK119Ub and H2BK120Ub as well as total histones H2A and H2B from isolated micronuclei (MN) and primary nuclei (PN) of MDA-MB-231 cells, *right*, relative H2AK119Ub and H2BK120Ub normalized to total histones H2A and H2B, respectively, * *p* < 0.05, two-sided t-test, n = 4 biological replicates (H2AK119Ub), n = 3 biological replicates (H2BK120Ub), bars represent mean ± SD.

**Supplementary Figure 3:**
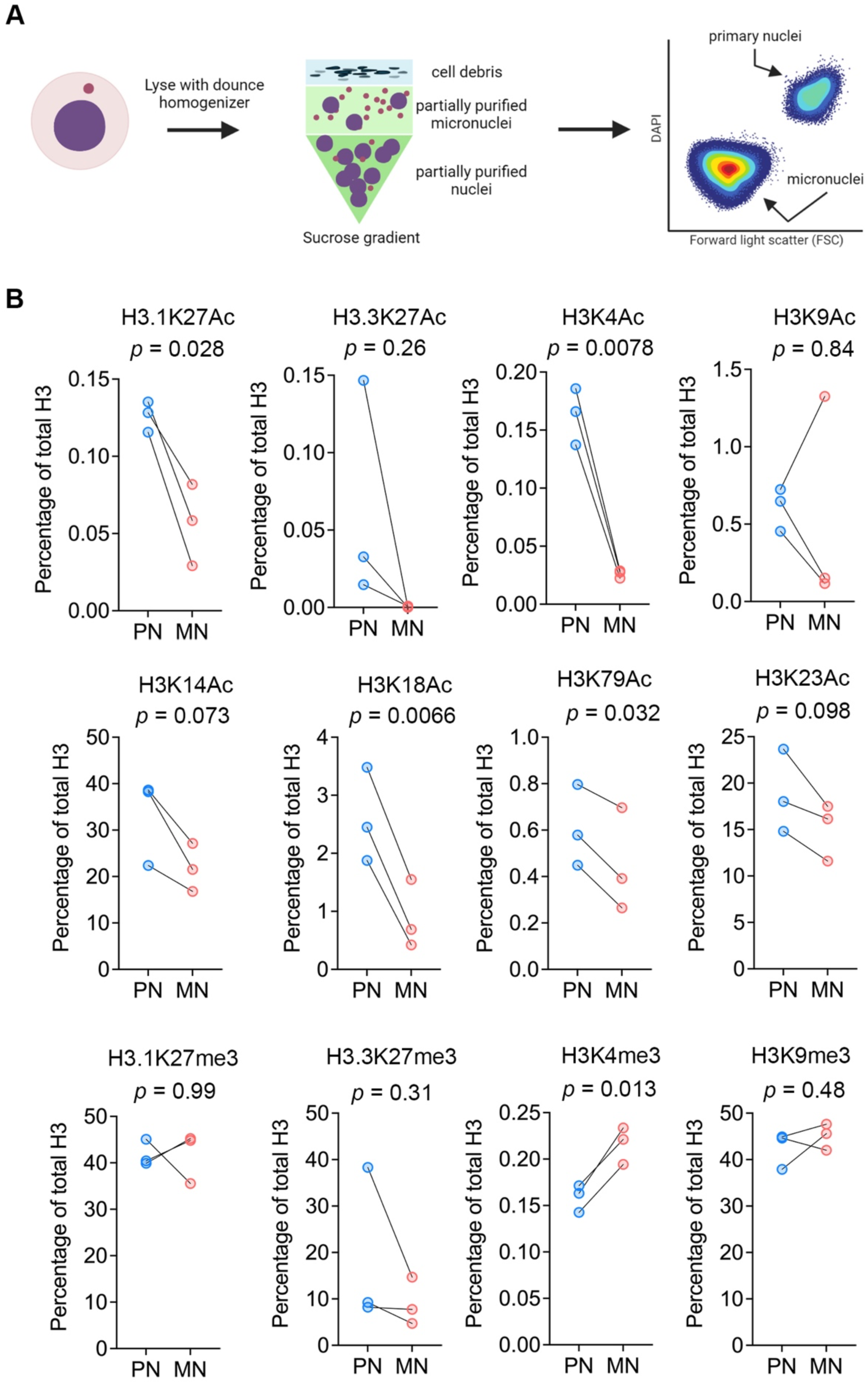
Mass spectrometry analysis of histone PTMs in primary nuclei and micronuclei. **A**, Micronuclei purification experiment schema. **B**, Abundance of various histone H3 PTMs relative to total histone H3 levels obtained from bottom-up histone mass spectrometry performed on isolated micronuclei (MN) and primary nuclei (PN), n = 3 biological replicates.

**Supplementary Figure 4:**
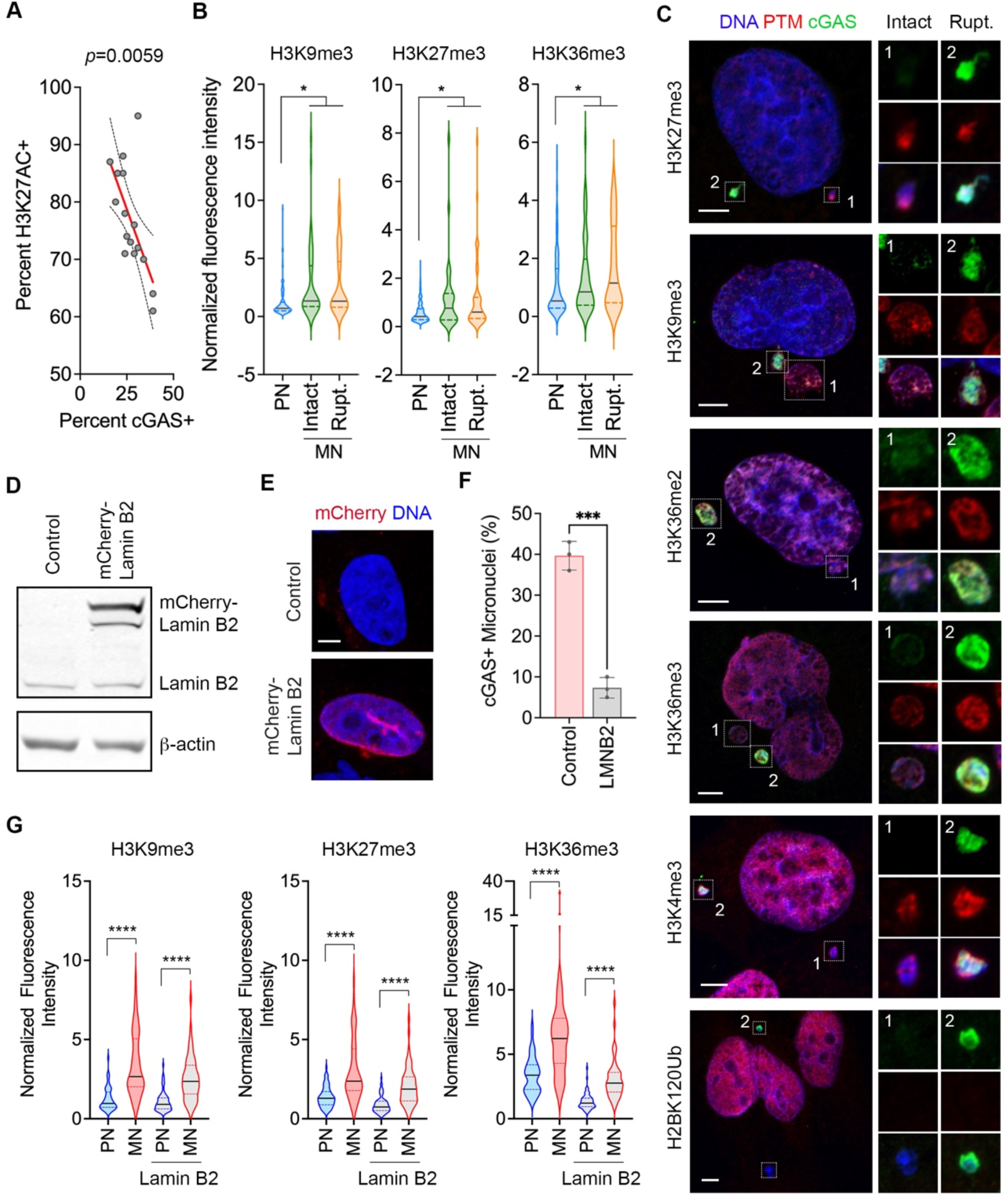
Impact of micronuclear rupture on histone PTMs. **A**, Sample-level correlation between percentages of micronuclei with H3K27Ac and cGAS staining in high grade serous ovarian cancer patient samples (n =16), 100 micronuclei counted / sample. **B**, Violin plot showing normalized fluorescence intensity of histone PTMs in MDA-MB-231 of micronuclei (MN) and primary nuclei (PN) in cells treated with vehicle control (DMSO) or GSK126, an EZH2 inhibitor (EZH2i); ns not significant, * *p* < 0.05, two-sided Mann-Whitney test), n > 50 cells, solid and dashed lines in the plot represent the median and quartiles, respectively. **C**, Representative immunofluorescence images of micronucleated MDA-MB-231 cells stained for DNA (blue), histone PTMs (red), and cGAS (green), scale bars 10 μm. **D**, Immunoblotting image of control and mCherry-LaminB2 overexpressing MDA-MB-231 cells with β-actin as loading control. **E**, Representative immunofluorescence image of control and mCherry-LaminB2 overexpressing MDA-MB-231 cells stained with DNA (blue), scale bar 10 μm. **F**, Percentage of micronuclei with cGAS staining in control and mCherry-LaminB2 overexpressing MDA-MB-231 cells (LMNB2); *** *p* < 0.001, two-sided t-test, n = 3 biological replicates, bars represent mean ± SD. **G**, Violin plot showing normalized fluorescence intensity of histone PTMs in primary nuclei and micronuclei of control and mCherry-Lamin B2 (Lamin B2) overexpressing MDA-MB-231 cells, n > 50 cells; **** *p* < 0.0001, two-sided Mann-Whitney test, solid and dashed lines in the plot represent the median and quartiles, respectively

**Supplementary Figure 5:**
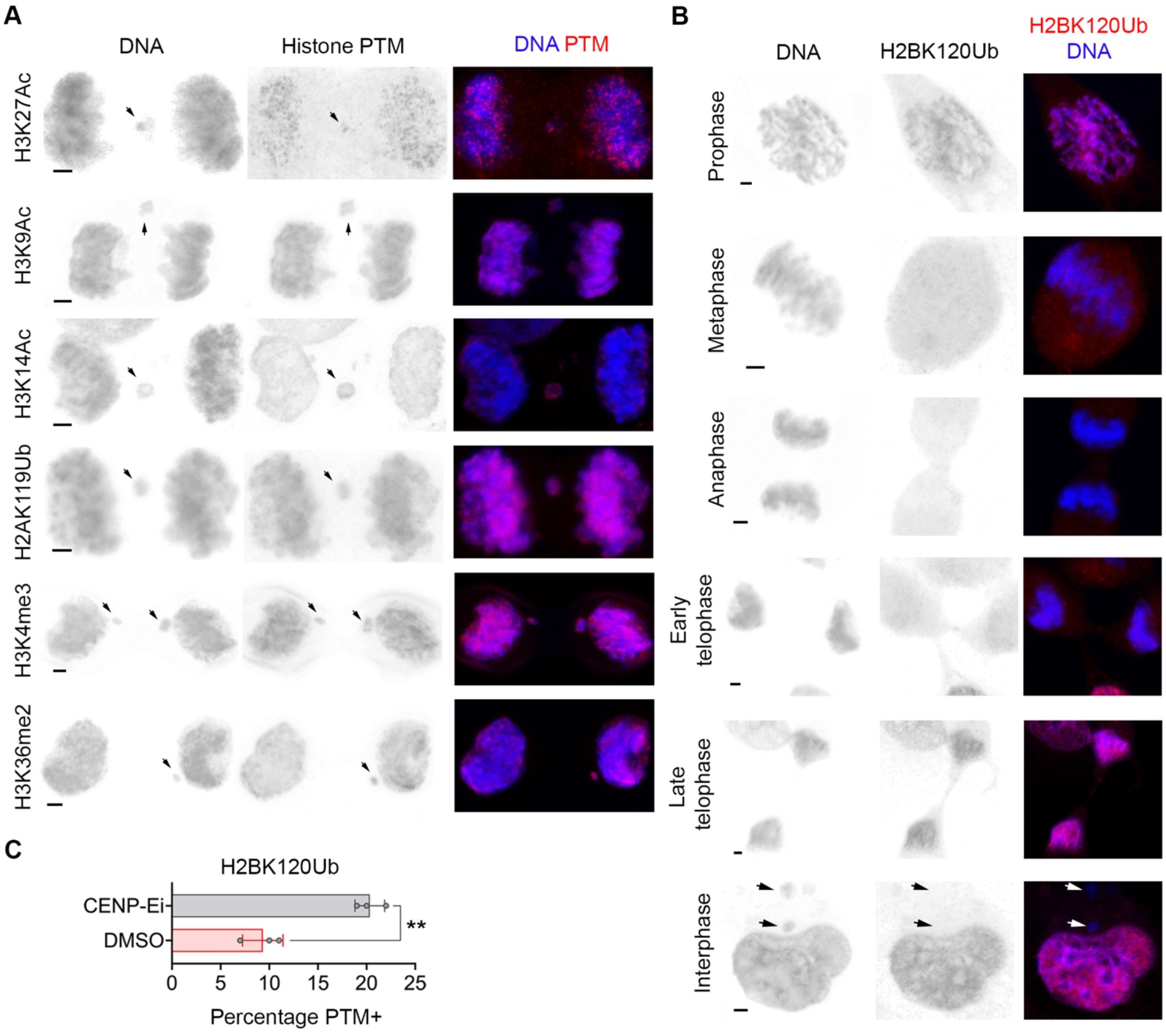
Mitotic errors are associated with abnormalities in histone PTMs. **A**, Representative immunofluorescence image of MDA-MB-231 cells with anaphase lagging chromosome (arrows) stained for DNA (blue) and histone PTMs (red). Scale bars 2 μm. **B**, Immunofluorescence images of MDA-MB-231 cells at different stages of the cell cycle, stained for DNA (blue) and H2BK120Ub (red), arrows denote micronuclei, scale bars 2 μm. **C**, Percentage of micronuclei staining for H2BK120Ub vehicle (DMSO)-treated or CENP-E inhibitor (CENP-Ei, GSK923295)-treated MDA-MB-231 cells; ** *p* < 0.01, two-sided t-test, n = 3 biological triplicates, bars represent mean ± SD.

**Supplementary Figure 6:**
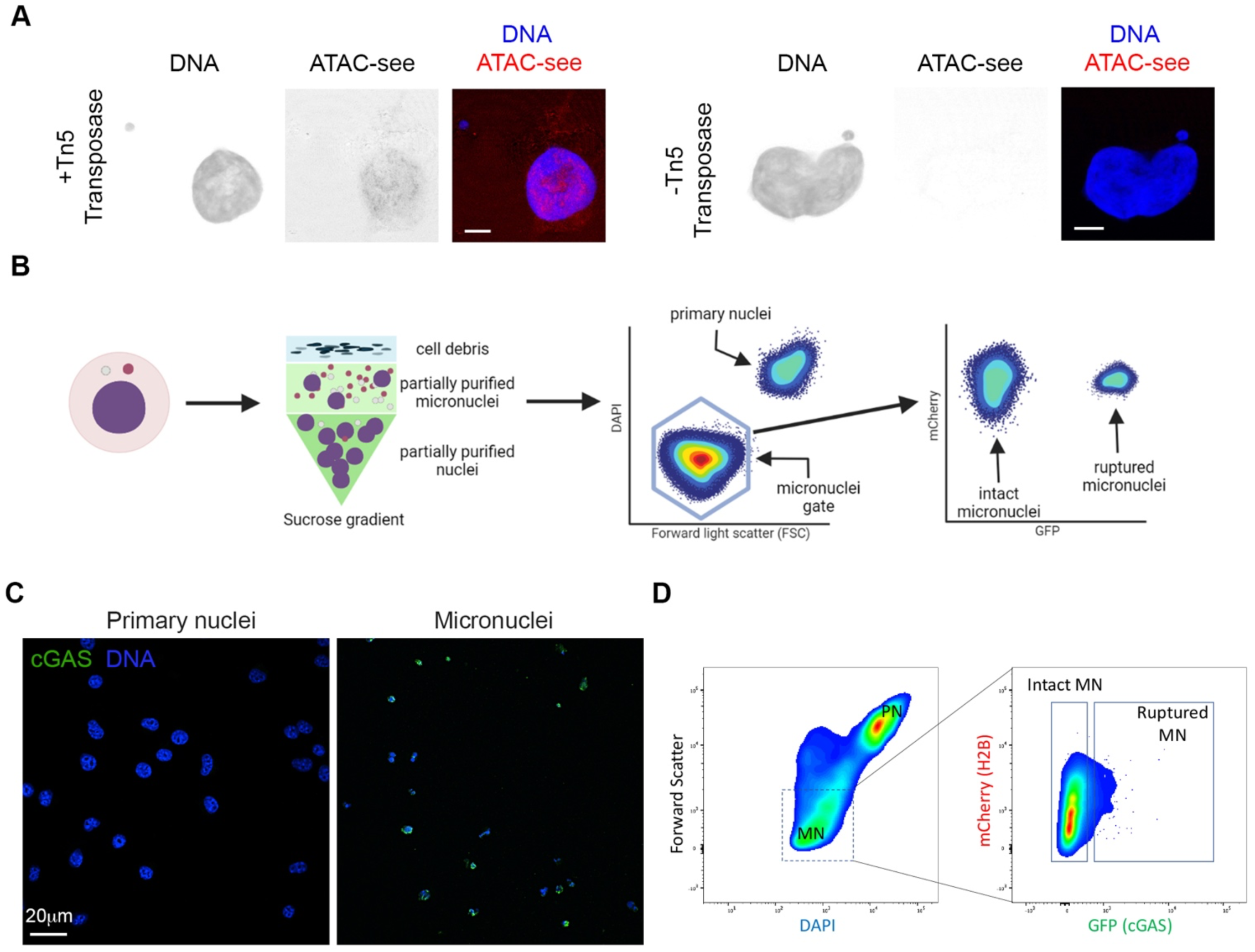
Micronuclei purification schema. **A**, Representative fluorescence images showing ATAC-see signal in MDA-MB-231 cells treated with (left) or without (right) fluorophore-tagged adaptors-loaded Tn5 transposase, scale bard 5 μm. **B**, Experimental schematic for the isolation of primary nuclei, intact micronuclei, and ruptured micronuclei. **C**, Representative immunofluorescence image from isolates of primary nuclei and micronuclei stained for DNA (blue) and cGAS (green), scale bar 20 μm. **D**, Density area plots of a representative micronuclei isolation flow cytometry experiment from 4T1 cells expressing H2B-mCherry and GFP-cGAS, right panel is the subset of the gate applied on the left panel (dashed box).

**Supplementary Figure 7:**
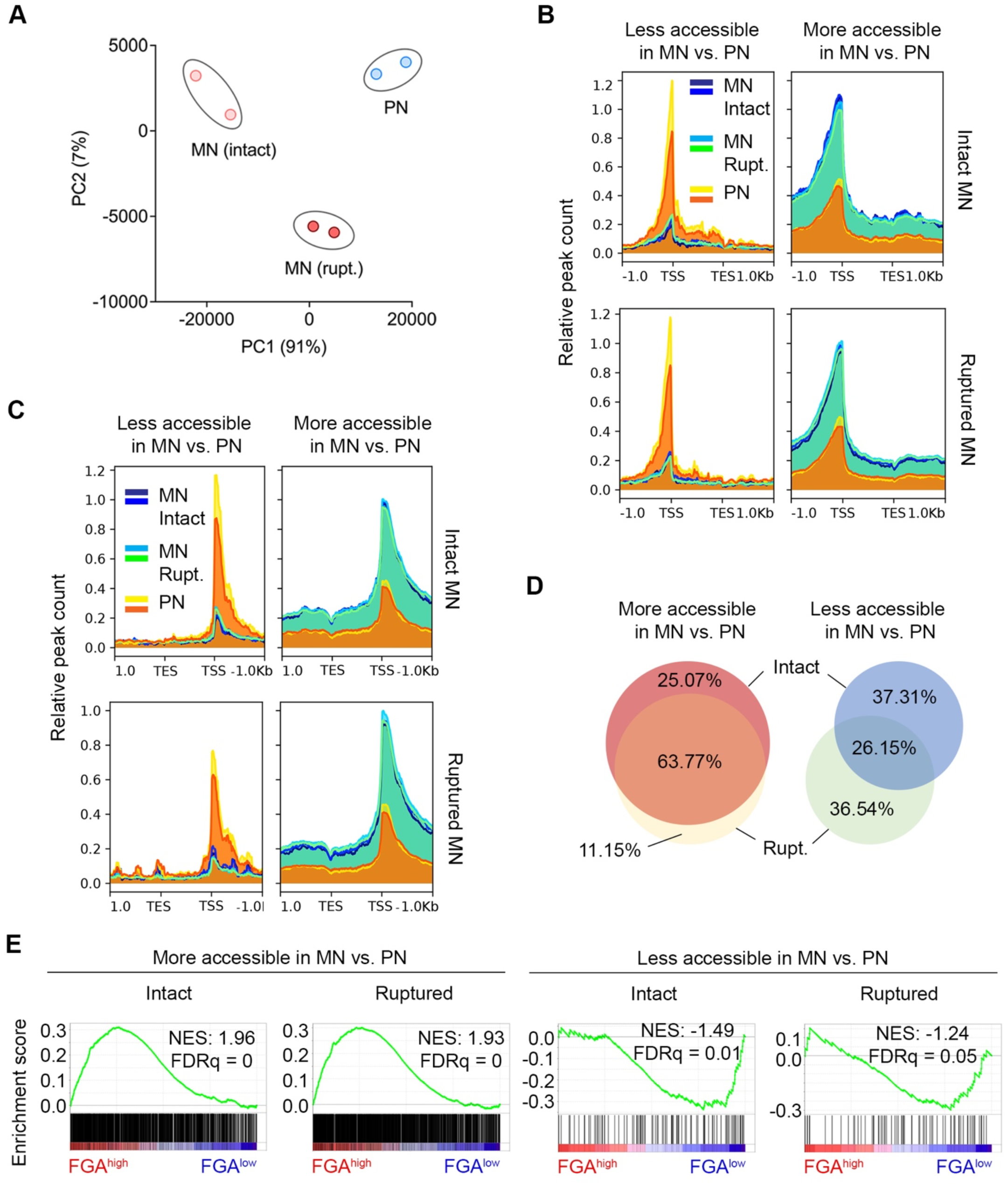
Chromatin accessibility patterns in micronuclei. **A**, Principal component analysis (PCA) plot of micronuclei (MN) and primary nuclei (PN) based on peak reads from ATAC-seq, n = 2 biological replicates. **B-C**, Density plot showing ATAC-Seq peak counts of differentially accessible positive (b) and negative (c)-strand genes in intact and ruptured micronuclei. **D**, Venn diagram representing the overlaps of in promoter accessibilities between intact and ruptured (Rupt.) micronuclei (MN) each relative to primary nuclei (PN). **E**, Enrichment plots of genes whose promoters are more (*left*) or less (*right*) enriched in micronuclei compared with primary nuclei comparing human breast tumors belonging to the top (FGA^high^) or bottom (FGA^low^) tertile of fraction genome altered according in the TCGA.

**Supplementary Figure 8:**
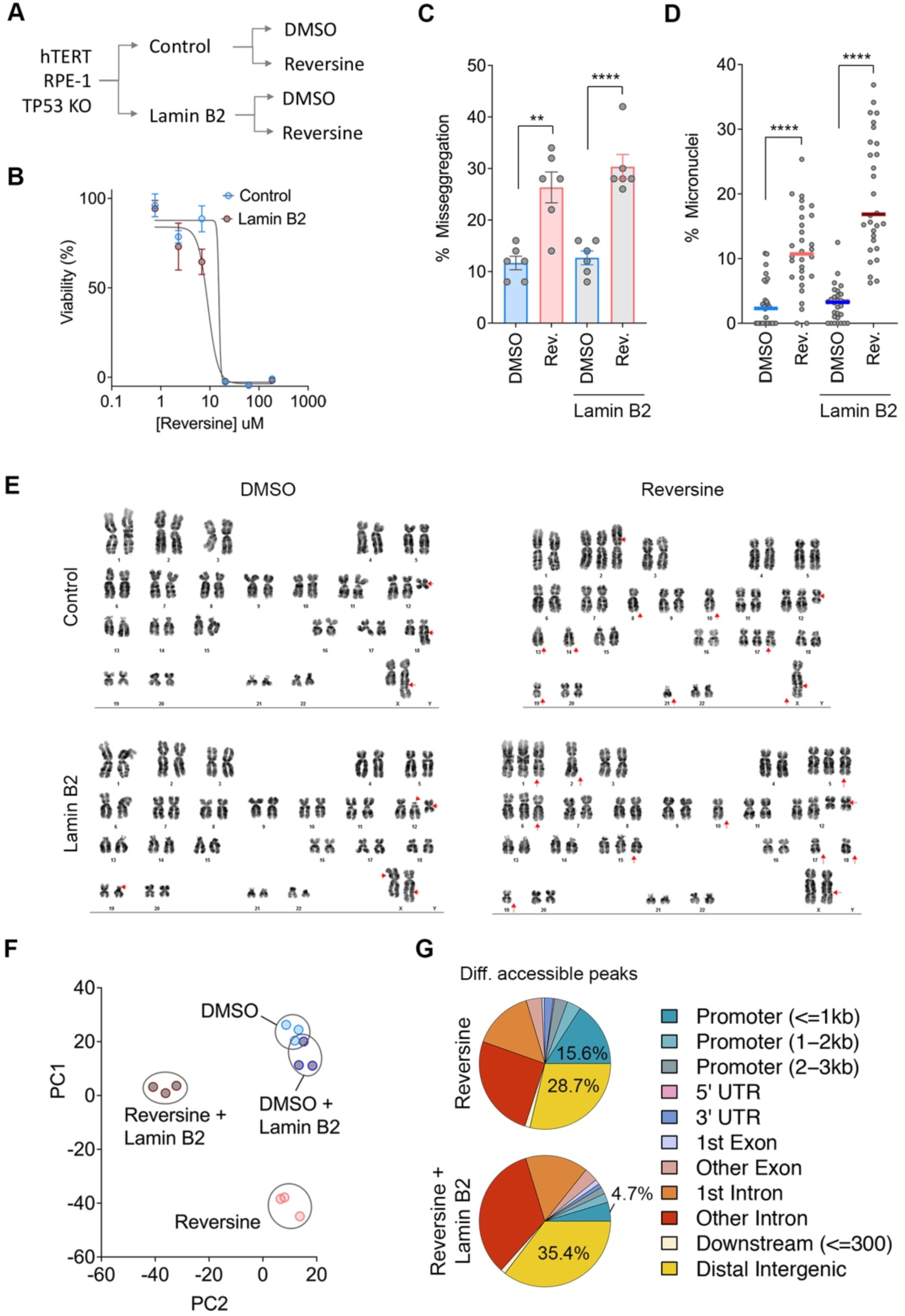
The generation of isogenic hTERT RPE-1 cell lines with different CIN rates. **A**, Experimental schema depicting the generation of isogenic hTERT RPE-1 cells with different CIN rates. **B**, IC50 curves of reversine-treated control and Lamin B2 overexpressing RPE-1 cells, points represent mean ± SD. **C**, Percentage of anaphase cells with chromosome mis-segregation in control or lamin B2 overexpressing RPE-1 cells long-term-treated vehicle (DMSO) or reversine (Rev.); n = 300 cells per condition, 6 biological replicates, *** *p* < 0.001, two-sided t-test, bars represent mean ± SD. **D**, Percentage of micronuclei relative to primary nuclei in control or lamin B2 overexpressing RPE-1 cells long-term-treated with vehicle (DMSO) or reversine (Rev.); n = 30 fields of view, 3 biological replicates, **** *p* < 0.0001, two-sided t-test, bars represent median. **E**, Conventional karyotypes of control or lamin B2 overexpressing RPE-1 cells long-term-treated with vehicle (DMSO) or reversine, red arrows denote chromosomal abnormalities. **F**, Principal component analysis (PCA) plot of TP53 KO hTERT-RPE1 cells based on the RNA expression of genes whose promoters exhibit differential accessibility in micronuclei compared to primary nuclei, n = 3 biological triplicates. **G**, Pie charts showing genomic regions of differentially accessible peaks that enriched (top panel) or depleted (bottom panel) in long-term reversine-treated compared with Lamin B2 overexpressing cells.

**Supplementary Figure 9.**
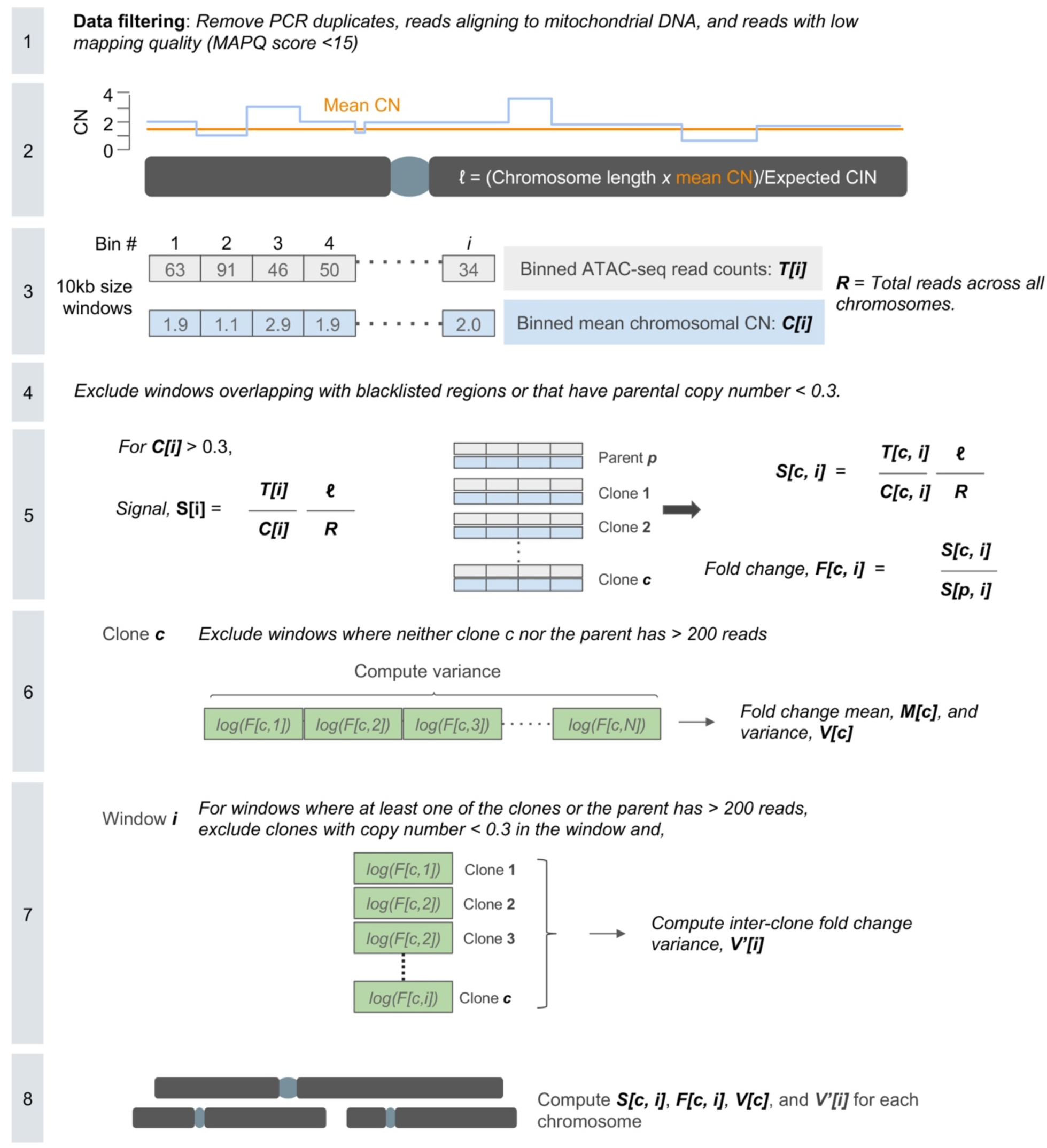
Experimental schematic for DLD-1 ATAC-Seq peaks normalization to copy number. More details can be found in the methods section “ATAC-Seq normalization for DLD-1 cells”.

**Supplementary Figure 10.**
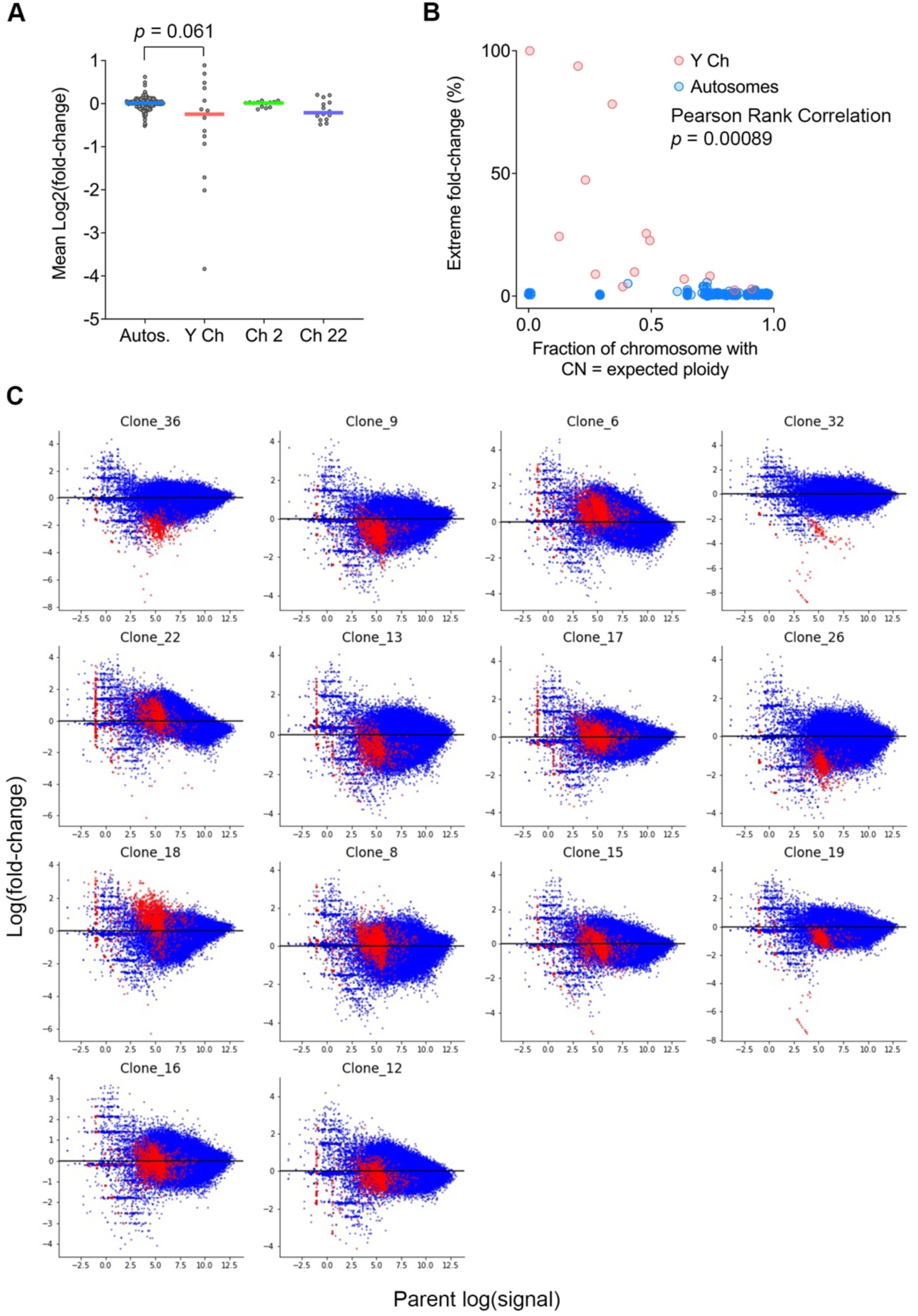
Chromosome missegregation and micronucleation promotes epigenetic dysregulation. **A**, Inter-clonal comparison of fold change of ATAC-seq peaks in autosomes, and the Y chromosomes in 14 DLD-1 single clones isolated from CEN-SELECT system. Chromosomes 2 and 22 were randomly selected as autosome representative, bars represent median, statistical significance tested using two-sided Mann-Whitney test. **B**, Percentage of autosomes or the Y chromosomes with extreme signal fold change in accessibility in the clones compared to the parental cell line as a function of chromosomal fraction with copy number (CN) equal to expected ploidy. Pearson Rank Correlation statistic comparing the Y chromosome values to all other chromosomes. **C**, Fold change distribution where y-axis represents the value of each clone’s fold change of the genomic window compared to the parental cell line, while x-axis represents the value of the parental cells’ log signal in the same window. Red dots = Y chromosomes, blue dots = autosomes.

## SUPPLEMENTARY DATA

### Supplementary Methods

#### Chromosome-based Normalization

In this section, we describe an alternative method for normalizing read counts. Instead of using the total ATAC-seq reads across all chromosomes, we normalize the reads in each genomic segment by the total ATAC-seq reads in just the chromosome containing the segment. The main motivation for using this method comes from the irregular distribution of total ATAC-seq reads per chromosome between the clones (**Supplementary Data Fig. 1**).

Using this method, the normalized signal of clone *c* in window *i* with total ATAC-seq reads *T*[*i*] and copy number *C*[*i*] was calculated as

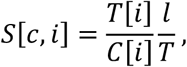

where *T* is the total ATAC-seq reads in the chromosome containing window *i* ; *l* is the ‘effective length’ of the chromosome and was calculated as

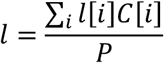

where *P* is the expected ploidy (P=1 for chromosome Y and P=2 for autosomes), and *l*[*i*], *C*[*i*] denote the length and copy number of window *i* respectively.

The signal fold-change of a clone *c* in window *i* was calculated as

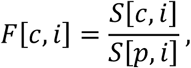

where *p* denotes the parental clone.

Similar to the genome-based normalization method (**Supplementary Fig. 9**), we calculated the normalized signal and fold change values for 10kb genomic segments and assessed inter-clonal and intra-clonal variance in accessibility. We observed that clones show a wider range of average fold-changes (**Supplementary Data Fig. 2A**) in Y chromosome, but not in autosomes. We also observed a higher variance in both intra-clonal and inter-clonal (**Supplementary Data Fig. 2B-C**) accessibility in Y chromosome. In addition, we see a correlation between the % of extreme fold-change values and the fraction of chromosome with copy number equal to expected ploidy in case of Y chromosome (**Supplementary Data Fig. 2D**).

## Supplementary Data Figures

**Supplementary Data Figure 1.**
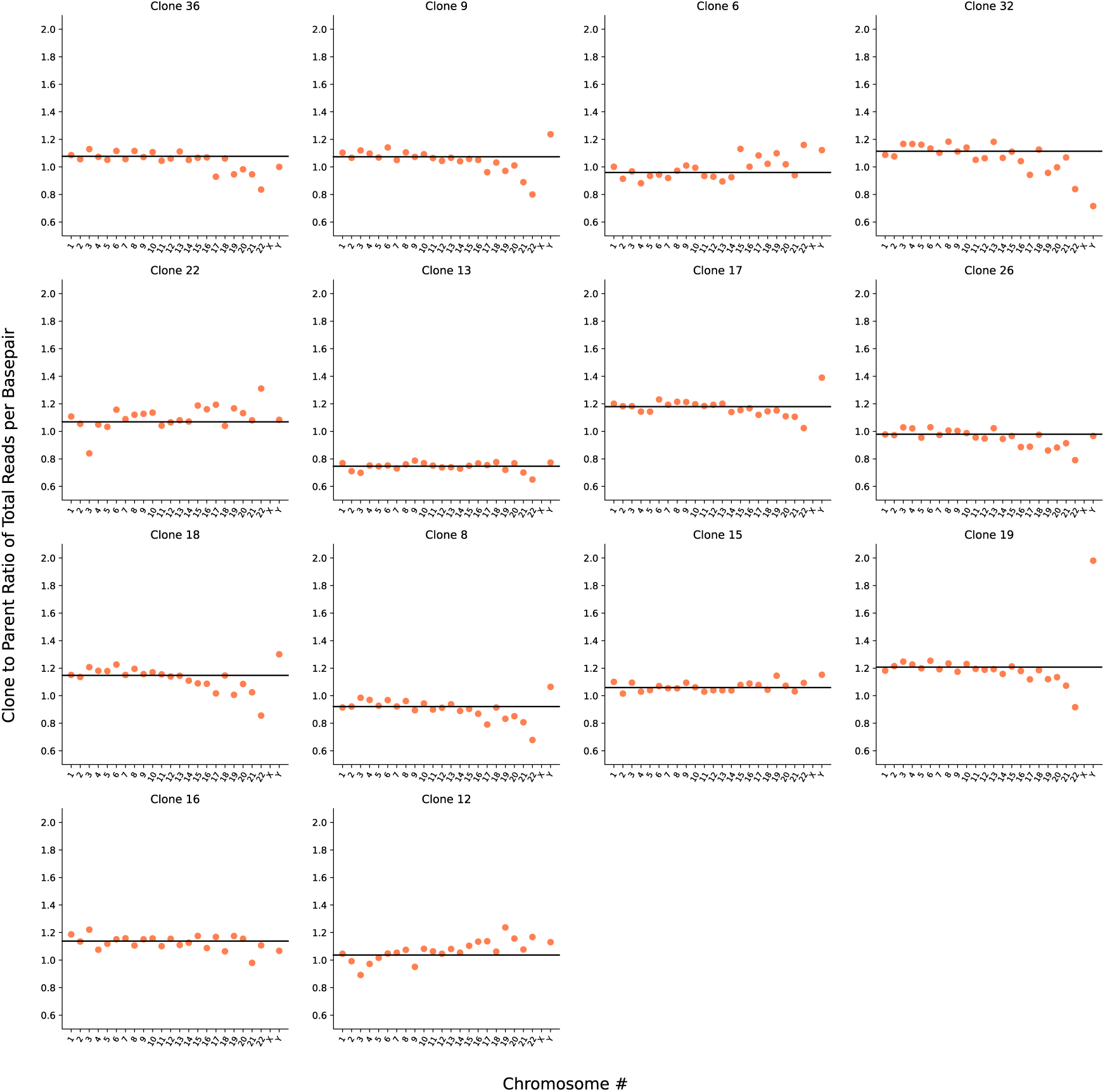
Irregular distribution of total ATAC-seq reads per base pair between clones. x-axis represents chromosome number and y-axis represents the ratio of total ATAC-seq reads per base pair in clone and parent. Dots show the ratio computed using total ATAC-seq reads in corresponding chromosome only. Horizontal black line represents the ratio computed using total ATAC-seq reads across all chromosomes.

**Supplementary Data Figure 2.**
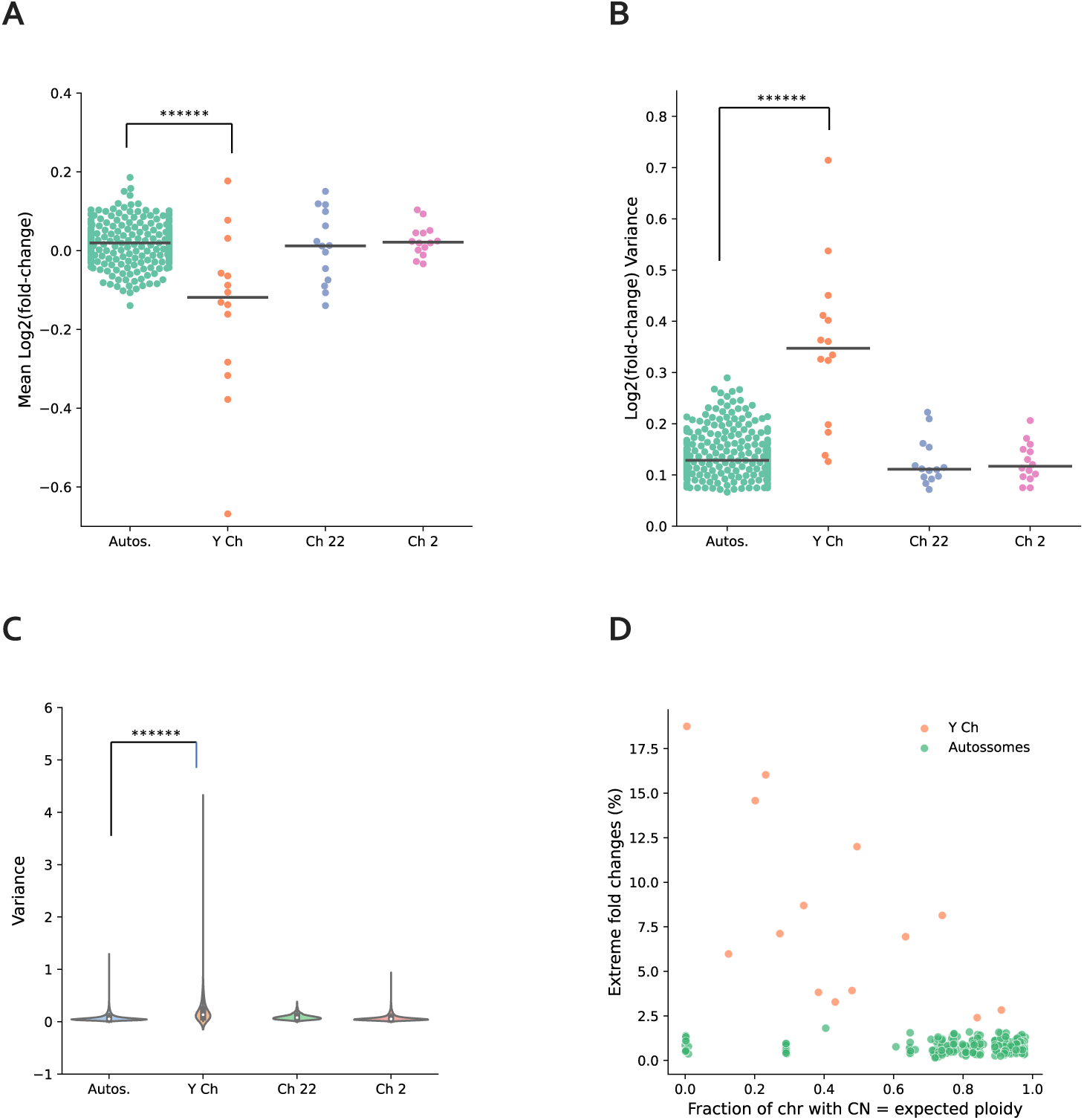
Increased epigenetic dysregulation in missegregated Y chromosome. **A** Comparison of average fold-changes across clones. Chromosomes 2 and 22 were randomly selected as autosome representative. Autos = autosomes, Y ch = Y chromosome, Ch 2 = chromosome 22, Ch 22 = chromosome 22. Bar represents median. ****** *p* < 0.000001, two-sided Mann-Whitney test. **B**, Inter-clonal variance of fold-changes. ****** *p* < 0.000001, two-sided Mann-Whitney test. **C**, Intra-clonal variance of fold-changes. ****** *p* < 0.000001, two-sided Mann-Whitney test. **D**, Extreme signal fold changes as a function of fraction of chromosomes with copy number (CN) equal to expected ploidy in Y chromosome and autosomes. *p* = 0.0186, Pearson rank correlation.

**Supplementary Table 1:** Antibodies used in immunofluorescence

| Antibody Against | Company | Catalog Number | Dilution |
| --- | --- | --- | --- |
| H3K4Me3 | Abcam | ab8580 | 1/1000 |
| H3K9Me3 | Abcam | ab8898 | 1/500 |
| H3K14Ac | Abcam | 52946 | 1/250 |
| H3K27Me3 | Active Motif | 61017 | 1/1000 |
| H3K27Ac | Abcam | ab4729 | 1/1000 |
| H3K36Me2 | Cell Signaling Technology | C75H12 | 1/200 |
| H3K36Me3 | Abcam | ab9050 | 1/1000 |
| H2AK119Ub | Cell Signaling Technology | 8240 | 1/1600 |
| H2BK120Ub | Cell Signaling Technology | 5546S | 1/1600 |
| cGAS (human) | LSBio | LS-C757990 | 1/1000 |
| cGAS (mouse) | Cell Signaling Technology | 31659S | 1/1000 |

**Supplementary Table 2:**
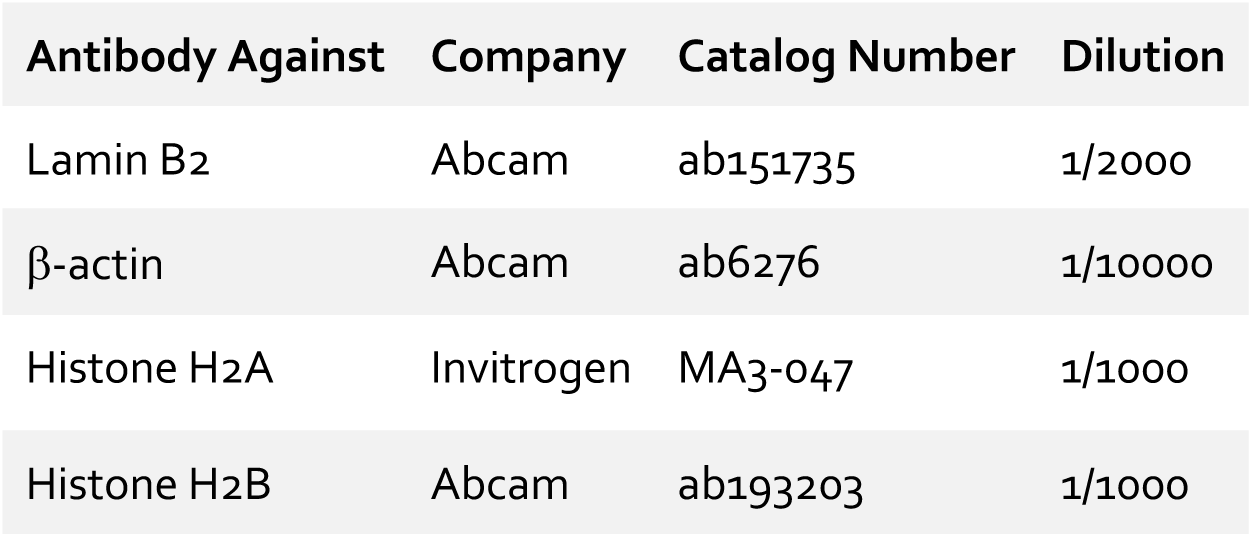
Antibodies used in immunoblotting

## REFERENCES

1. Jones, P.A. and S.B. Baylin, The epigenomics of cancer. Cell, 2007. 128(4): p. 683–92.

2. Bakhoum, S.F. and L.C. Cantley, The Multifaceted Role of Chromosomal Instability in Cancer and Its Microenvironment. Cell, 2018. 174(6): p. 1347–1360.

3. Timp, W. and A.P. Feinberg, Cancer as a dysregulated epigenome allowing cellular growth advantage at the expense of the host. Nat Rev Cancer, 2013. 13(7): p. 497–510.

4. Watkins, T.B.K., et al., Pervasive chromosomal instability and karyotype order in tumour evolution. Nature, 2020.

5. Kato, H. and A.A. Sandberg, Chromosome pulverization in human cells with micronuclei. J Natl Cancer Inst, 1968. 40(1): p. 165–79.

6. Crasta, K., et al., DNA breaks and chromosome pulverization from errors in mitosis. Nature, 2012. 482(7383): p. 53–8.

7. Hatch, E.M., et al., Catastrophic nuclear envelope collapse in cancer cell micronuclei. Cell, 2013. 154(1): p. 47–60.

8. Lengauer, C., K.W. Kinzler, and B. Vogelstein, Genetic instability in colorectal cancers. Nature, 1997. 386(6625): p. 623–7.

9. Lengauer, C., K.W. Kinzler, and B. Vogelstein, Genetic instabilities in human cancers. Nature, 1998. 396(6712): p. 643–9.

10. Burrell, R.A., et al., The causes and consequences of genetic heterogeneity in cancer evolution. Nature, 2013. 501(7467): p. 338–45.

11. Bakhoum, S.F., et al., Chromosomal instability drives metastasis through a cytosolic DNA response. Nature, 2018. 553(7689): p. 467–472.

12. Lee, A.J., et al., Chromosomal instability confers intrinsic multidrug resistance. Cancer Res, 2011. 71(5): p. 1858–70.

13. Davoli, T., et al., Tumor aneuploidy correlates with markers of immune evasion and with reduced response to immunotherapy. Science, 2017. 355(6322).

14. McGranahan, N., et al., Allele-Specific HLA Loss and Immune Escape in Lung Cancer Evolution. Cell, 2017. 171(6): p. 1259–1271 e11.

15. Thompson, S.L. and D.A. Compton, Examining the link between chromosomal instability and aneuploidy in human cells. J Cell Biol, 2008. 180(4): p. 665–72.

16. Mackenzie, K.J., et al., cGAS surveillance of micronuclei links genome instability to innate immunity. Nature, 2017. 548(7668): p. 461–465.

17. Harding, S.M., et al., Mitotic progression following DNA damage enables pattern recognition within micronuclei. Nature, 2017. 548(7668): p. 466–470.

18. Ly, P., et al., Chromosome segregation errors generate a diverse spectrum of simple and complex genomic rearrangements. Nat Genet, 2019. 51(4): p. 705–715.

19. Zhang, C.Z., et al., Chromothripsis from DNA damage in micronuclei. Nature, 2015. 522(7555): p. 179–84.

20. Mohr, L., et al., ER-directed TREX1 limits cGAS activation at micronuclei. Mol Cell, 2021. 81(4): p. 724–738 e9.

21. Yang, H., et al., cGAS is essential for cellular senescence. Proc Natl Acad Sci U S A, 2017. 114(23): p. E4612–E4620.

22. Wang, F. and J.M. Higgins, Histone modifications and mitosis: countermarks, landmarks, and bookmarks. Trends Cell Biol, 2013. 23(4): p. 175–84.

23. Duan, Y., et al., Ubiquitin ligase RNF20/40 facilitates spindle assembly and promotes breast carcinogenesis through stabilizing motor protein Eg5. Nat Commun, 2016. 7: p. 12648.

24. Kapoor, T.M., et al., Chromosomes can congress to the metaphase plate before biorientation. Science, 2006. 311(5759): p. 388–91.

25. Bennett, A., et al., Cenp-E inhibitor GSK923295: Novel synthetic route and use as a tool to generate aneuploidy. Oncotarget, 2015. 6(25): p. 20921–32.

26. Abdollahi, E., G. Taucher-Scholz, and B. Jakob, Application of fluorescence lifetime imaging microscopy of DNA binding dyes to assess radiation-induced chromatin compaction changes. Int J Mol Sci, 2018. 19(8).

27. Chen, X., et al., ATAC-see reveals the accessible genome by transposase-mediated imaging and sequencing. Nat Methods, 2016. 13(12): p. 1013–1020.

28. Santaguida, S., et al., Dissecting the role of MPS1 in chromosome biorientation and the spindle checkpoint through the small molecule inhibitor reversine. J Cell Biol, 2010. 190(1): p. 73–87.

29. Ly, P., et al., Selective Y centromere inactivation triggers chromosome shattering in micronuclei and repair by non-homologous end joining. Nat Cell Biol, 2017. 19(1): p. 68–75.

30. Shoshani, O., et al., Chromothripsis drives the evolution of gene amplification in cancer. Nature, 2021. 591(7848): p. 137–141.

31. McDonald, O.G., et al., Epigenomic reprogramming during pancreatic cancer progression links anabolic glucose metabolism to distant metastasis. Nat Genet, 2017. 49(3): p. 367–376.

32. Hoadley, K.A., et al., Cell-of-Origin Patterns Dominate the Molecular Classification of 10,000 Tumors from 33 Types of Cancer. Cell, 2018. 173(2): p. 291–304 e6.

33. Malta, T.M., et al., Machine Learning Identifies Stemness Features Associated with Oncogenic Dedifferentiation. Cell, 2018. 173(2): p. 338–354 e15.

34. Chan, H.L., et al., Polycomb complexes associate with enhancers and promote oncogenic transcriptional programs in cancer through multiple mechanisms. Nat Commun, 2018. 9(1): p. 3377.

35. Comet, I., et al., Maintaining cell identity: PRC2-mediated regulation of transcription and cancer. Nat Rev Cancer, 2016. 16(12): p. 803–810.

36. Kadoch, C. and G.R. Crabtree, Mammalian SWI/SNF chromatin remodeling complexes and cancer: Mechanistic insights gained from human genomics. Sci Adv, 2015. 1(5): p. e1500447.

37. Hose, J., et al., Dosage compensation can buffer copy-number variation in wild yeast. Elife, 2015. 4.

38. Huang, Y., L. Gu, and G.M. Li, H3K36me3-mediated mismatch repair preferentially protects actively transcribed genes from mutation. J Biol Chem, 2018. 293(20): p. 7811–7823.

39. Pai, C.C., et al., A histone H3K36 chromatin switch coordinates DNA double-strand break repair pathway choice. Nat Commun, 2014. 5: p. 4091.

40. Wu, S., et al., Circular ecDNA promotes accessible chromatin and high oncogene expression. Nature, 2019. 575(7784): p. 699–703.

41. Turner, K.M., et al., Extrachromosomal oncogene amplification drives tumour evolution and genetic heterogeneity. Nature, 2017. 543(7643): p. 122–125.

42. Allis, C.D., et al., Deposition-related histone acetylation in micronuclei of conjugating Tetrahymena. Proc Natl Acad Sci U S A, 1985. 82(23): p. 8048–52.

43. Toufektchan, E. and J. Maciejowski, Purification of micronuclei from cultured cells by flow cytometry. STAR Protoc, 2021. 2(1): p. 100378.

44. Cerami, E., et al., The cBio cancer genomics portal: an open platform for exploring multidimensional cancer genomics data. Cancer Discov, 2012. 2(5): p. 401–4.

45. Buenrostro, J.D., et al., ATAC-seq: A Method for Assaying Chromatin Accessibility Genome-Wide. Curr Protoc Mol Biol, 2015. 109: p. 21 29 1–21 29 9.

46. Langmead, B. and S.L. Salzberg, Fast gapped-read alignment with Bowtie 2. Nat Methods, 2012. 9(4): p. 357–9.

47. Sidoli, S., et al., Complete Workflow for Analysis of Histone Post-translational Modifications Using Bottom-up Mass Spectrometry: From Histone Extraction to Data Analysis. J Vis Exp, 2016(111).

48. Yuan, Z.F., et al., EpiProfile 2.0: A Computational Platform for Processing Epi-Proteomics Mass Spectrometry Data. J Proteome Res, 2018. 17(7): p. 2533–2541.

